# The submergence-induced drastic morphological plasticity of root in the amphibious plant *Callitriche palustris*

**DOI:** 10.64898/2026.04.08.716617

**Authors:** Tomo Sato, Yuki Doll, Mikiko Kojima, Yumiko Takebayashi, Jun Takeuchi, Yasushi Todoroki, Hitoshi Sakakibara, Hiroyuki Koga, Hirokazu Tsukaya

**Affiliations:** Graduate School of Science, The University of Tokyo, 7-3-1, Hongo, Bunkyo-ku, Tokyo, 113-0033, Japan; Division of Biological Science, Graduate School of Science and Technology, Nara Institute of Science and Technology, Ikoma, Nara, 8916-5, Japan; Mass Spectrometry and Microscopy Unit, RIKEN Center for Sustainable Resource Science (CSRS), Suehiro-cho, Tsurumi-ku, Yokohama, Kanagawa, 230-0045, Japan; Faculty of Agriculture, Shizuoka University, Ohya, Suruga-ku, Shizuoka 422-8529, Japan; Graduate School of Bioagricultural Sciences, Nagoya University, Nagoya, Aichi, 464-8601, Japan

## Abstract

Amphibious plants can thrive in both terrestrial and submerged environments, which are fundamentally distinct. Although morphological plasticity of leaf known as heterophylly has been well investigated, the morphological plasticity of root in amphibious plants remains poorly understood. In this study, we discovered that an amphibious plant *Callitriche palustris* (Plantaginaceae), which has significant heterophylly, has a remarkable morphological plasticity also in root in response to submergence. This species develops thin roots with abundant root hairs, fewer cortical and epidermal cells, and smaller aerenchyma in the terrestrial condition. On the other hand, it develops thicker roots with few root hairs, more cortical and epidermal cells, and larger aerenchyma in the submerged condition. We call this morphological plasticity of root as “heterorhizy”. Phytohormone perturbation experiments revealed that abscisic acid (ABA) and gibberellin regulate root hair development and root cell division respectively. We also found the possibility that heterorhizy was acquired in the genus *Callitriche*. Additionally, a similar form of root hair plasticity was also observed in the phylogenetically distinct amphibious species *Ludwigia arcuata* (Onagraceae). Furthermore, the absence of root hair development underwater and the similar structure of aerenchyma to *C. palustris* were broadly seen across diverse aquatic plants. This study provides new insights into the root morphological responses to submerged environments in aquatic plants.

## Introduction

More than 450 million years ago, land plants evolved from aquatic algae, proceeded to the terrestrial condition, and diverged into hundreds of thousands of species (Delwiche & Cooper, 2015; Lughadha et al. 2016). Most of the current land plants are well adapted to the terrestrial condition, and submergence is a fatal stress for many land plants (Bailey-Serres & Voesenek, 2008). Nevertheless, numerous lineages of plants have repeatedly adapted to life in the water. Such plant species are referred to as “secondary aquatic plants” (Arber, 1920; Cook, 1999). In this article, secondary aquatic plants are referred to simply as “aquatic plants”. Among aquatic plants, those capable of surviving in both terrestrial and submerged environments are known as “amphibious plants" (Arber, 1920; Koga et al. 2024; Sculthorpe, 1967). Aquatic lifestyles have evolved independently more than one hundred times across land plants, representing a remarkable example of convergent evolution (Cook, 1999; Guo et al. 2025).

Submergence imposes multiple physiological constraints on plants, among which oxygen limitation is particularly detrimental to roots (Bailey-Serres & Voesenek, 2008; Fukao et al. 2019). Oxygen deficiency caused by submergence severely impairs root respiration, leading to hypoxia that threatens root survival (Colmer, 2003; Colmer et al. 2020). To cope with oxygen deficiency, many wetland plants form intercellular gas-exchange tissue in roots, known as aerenchyma (Schenk, 1889; Seago et al. 2005; Jung et al. 2008; Yamauchi et al. 2013; Takahashi et al. 2014). For example, Poaceae crop plants such as *Oryza sativa* and *Zea mays* form aerenchyma in the root cortical layers as an adaptive response to waterlogging (Yamauchi et al. 2013; Takahashi et al. 2014). Aerenchyma facilitates internal oxygen transport to root tips and contributes to the maintenance of root viability under low oxygen conditions (Colmer, 2003; Abiko et al. 2012; Yamauchi et al. 2018). There are typically two types of aerenchyma: one is lysigenous aerenchyma produced by cell death and the other is schizogenous aerenchyma produced by cell separation and differential cell division and/or cell expansion (Arber, 1920; Smirnoff & Crawford, 1983; Seago et al. 2005). Some studies revealed that phytohormones such as auxin and ethylene are important in the formation of lysigenous aerenchyma in Poaceae plants (Yamauchi et al. 2016, 2020). Also, a study reported the basic morphological characteristics of schizogenous aerenchyma of the wetland Brassicaceae plant *Cardamine amara* (Kudoh et al. 2025). Together, these studies have been focusing on root anatomical adaptations and their developmental regulation in wetland plants with roots embedded in waterlogged soils.

In contrast, amphibious plants often experience complete submergence and develop roots that grow freely in the water column, without contact with soil. Although some studies reported that root hairs are absent in roots growing freely in water in some aquatic plants (Cormack, 1937; Shannon, 1953; Clayton & Bagyaraj, 1984; Yin et al. 2025), these observations were mainly based on submerged plants or floating plants, and whether amphibious plants exhibit submergence-induced plasticity in root development, including root hair development, has not been systematically investigated.

Previous studies of amphibious plants have been focusing mainly on the leaf phenotypic plasticity. Many amphibious plants develop different shapes of leaves in response to the two conditions, which is called heterophylly (Arber, 1920; Sculthorpe, 1967; Nakayama et al. 2017; Li et al., 2019; Koga et al. 2024). In the terrestrial condition, many amphibious plants develop round or less serrated leaves, harboring many stomata, thick cell wall and cuticular layer on the epidermis. On the other hand, in the submerged condition, they develop linear or more serrated leaves, with less stomata, thin cell wall, and cuticular layer (Kuwabara et al. 2001; Nakayama et al. 2014; Li et al. 2017; Kim et al. 2018; Koga et al. 2020). These morphological changes help amphibious plants to adapt to the two distinct environmental conditions (Nielsen, 1993; Wells & Pigliucci, 2000; Mommer & Visser, 2005; Horiguchi et al. 2019). The molecular mechanisms of heterophylly were also investigated in several species, and previous research revealed that phytohormones such as abscisic acid (ABA), ethylene, and gibberellin (GA) are key factors of heterophylly and exogenous phytohormone treatment can change the leaf shape (Anderson, 1978; Goliber & Feldman, 1989; Kuwabara et al. 2003; Nakayama et al. 2017; Li et al. 2019; Koga et al. 2024; Sakamoto et al. 2024). These studies suggest that amphibious plants have similar aspects of regulatory mechanism in their heterophylly, despite the independent origins of amphibious lifestyles. Given this pronounced leaf plasticity, it is reasonable to hypothesize that amphibious plants may also exhibit environmentally induced plasticity in roots.

In this study, we investigated the root morphology under terrestrial and submerged conditions in an amphibious plant *Callitriche palustris* (Plantaginaceae). We previously demonstrated that this species has remarkable heterophylly and its regulatory mechanism by phytohormones such as ABA, ethylene, and GA (Koga et al. 2020, 2021). In addition, the genus *Callitriche* includes diverse lifestyles of species, including terrestrial, amphibious, and obligately submerged species (Ito et al. 2017; Doll et al. 2021), making it an ideal system for studying the evolution of aquatic and amphibious lifestyles. Here, using *C. palustris*, we examined the root morphology under terrestrial and submerged conditions and discovered drastic morphological plasticity on root hair development and internal structure between the two conditions, a form of root morphological plasticity that we term “heterorhizy.” We then examined the effect of phytohormone on this morphological plasticity and investigated its evolutionary process using relative *Callitriche* species. Finally, we observed a similar plasticity of root hair development in *Ludwigia arcuata* (Onagraceae), a phylogenetically distinct amphibious species, and similar morphological features in other aquatic plants across diverse lineages.

## Materials and methods

### Plant culture

A plant of *Callitriche palustris* was originally collected in Hakuba, Nagano, Japan. The plant material used in this study was an inbred line generated by 11 generations of self-fertilization, and multiple sister individuals were used for all experiments. The basic plant culture system used in this study was previously described (Koga et al. 2020). Briefly, the plants were asexually propagated by stem cutting and grown on 100 mL of solid half-strength Murashige and Skoog medium (FUJIFILM Wako Pure Chemical Corporation, Osaka, Japan) solidified with 0.3% (w/v) gellan gum (Wako, Osaka, Japan) (Fig. 1A). Plants were grown with a long-day (16 h light and 8 h dark) condition at 22℃ with a light intensity of 60 μmol m^−2^ s^−1^ for two weeks. As the submerged condition, 200 mL of sterile distilled water was poured on the medium.

**Figure 1.**
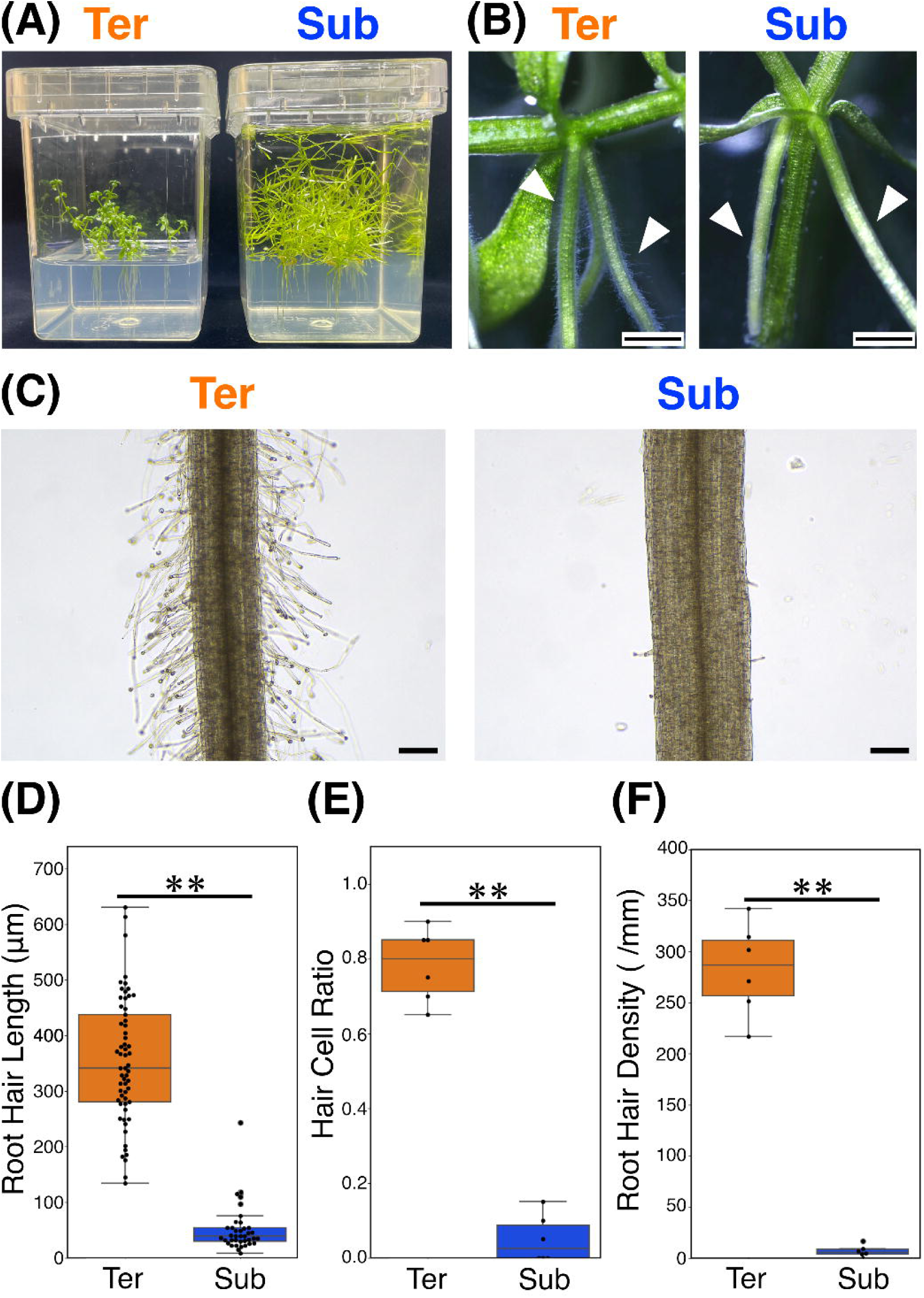
Morphological plasticity of root in *Callitriche palustris*. **(A)** Standard cultivation system used in this study. **(B)** Adventitious roots formed at stem nodes. Arrows indicate adventitious roots. **(C)** Morphological plasticity of root hair under terrestrial and submerged conditions. **(D)** Root hair length under the two conditions. **(E)** Proportion of root hair cells among all epidermal cells. **(F)** Root hair density under the two conditions. Asterisks indicate significant differences between conditions (∗p < 0.05; ∗∗p < 0.01; Welch’s t-test). Scale bars: **(B)** 1 mm; **(C)** 100 μm.

For the phytohormone treatments, a given concentration of phytohormones or the signaling inhibitors was dissolved from stock solution into both medium and water. Abscisic acid (ABA) (Tokyo Chemical Industry, Tokyo, Japan), gibberellin A3 (GA₃) (Tokyo Chemical Industry), and the ethylene precursor 1-Aminocyclopropane-1-carboxylic acid (ACC) (Tokyo Chemical Industry) were used as signaling activators, whereas the ABA antagonist PANMe, the gibberellin biosynthesis inhibitor Uniconazole P (Wako), and the ethylene signaling inhibitor silver nitrate (AgNO₃) (Wako) were used as signaling inhibitors (Rademacher, 2000; Takeuchi et al. 2018; Binder, 2020). Since the ABA inhibitor PANMe (Takeuchi et al. 2018) was dissolved in DMSO (Nacalai tesque, Kyoto, Japan) and GA_3_ was prepared as a stock solution in 100% (v/v) ethanol (Nacalai tesque), DMSO and ethanol were used as the corresponding control treatments. The working concentration for each phytohormone was defined as the highest dose that elicited phenotypic responses without severely inhibiting root development, based on preliminary experiments.

A simple, custom-built hydroponic culture system was established for this study (see Supplementary Fig. 2A). First, two pairs of half-cut glass slides were fixed with the cut corner of glass slides using a water unsolvable glue. This glass slide basis was put in the plant container and autoclaved. Also, 15 mL of thin solid medium was solidified into square shape with stainless mesh. After pouring approximately 100 mL of liquid MS medium on the glass unit, which contains the same component except gellan gum solidifier, solid medium was put on it.

Other *Callitriche* species were also collected in Japan, and *Ludwigia arcuata* was purchased from an aquarium shop. These species were also grown on sterile conditions using MS media or Gamborg’s B5 media (FUJIFILM Wako Pure Chemical Corporation), indicated in Supplementary table 1. Other aquatic plants were also collected in wild or commercially purchased (Supplementary table 2).

After cultivation, samples were fixed with a formalin-acetic acid-alcohol (FAA) fixative (2.5% (v/v) formalin (Wako), 2.5% acetic acid (v/v) (Nacalai tesque), and 45% (v/v) ethanol [v/v] in Milli-Q water) and stored at room temperature. All experiments were replicated at least two times.

### Measurement of root hair length and density

For the quantification of root hair traits, root segments were excised at 5 mm point from the root tip. Newly developed roots after cutting in solid medium without touching the bottom of plant container were chosen for analysis. The segments were stained with 1% (v/v) Calcofluor white (Sigma-Aldrich, Darmstadt, Germany) in ClearSee solution (Kurihara et al. 2015) and mounted on glass slides with the staining solution. Pictures were obtained with a confocal microscope (Fv10i, Olympus, Tokyo, Japan). Obtained images were stitched using Grid/Collection stitching plugin in ImageJ/Fiji (Preibisch et al. 2009; Schindelin et al. 2012). Root hair length and root hair density were manually measured from one half side of the root surface. Root hair density was calculated by the root hair number divided by observed root length (mm). To normalize the differences in epidermal cell numbers between the terrestrial and submerged conditions, the hair cell ratio was also used. This metric was defined as the proportion of root hair cells among 20 contiguous epidermal cells. Fiji/ImageJ software was used for image analysis (Schindelin et al. 2012, Ver. 2.14.0, US National Institutes of Health).

### Analysis of root anatomy

Root cross-sections were made from root segments excised at 10 mm point from the root tip. Segments were embedded in Technovit 7100 resin (Kulzer, Hanau, Germany). First, segments were pre-stained with 0.1% (w/v) Safranin O (Waldeck GmbH & Co., KG Division Chroma) in 50% ethanol, and gradually dehydrated with ethanol series for each 10 minutes. The segments were washed in a solution of 50% Technovit 7100 solution in ethanol twice and 100% solution. Next, the samples were stained with Neutral Red (Wako) in Technovit solution and then washed with the normal solution. Finally, the segments were immersed in 300 μL of polymerization solution (1/13 Technovit 7100 Hardener II in Technovit 7100) placed in the cut lid of a 1.5 mL tube (INA・OPTIKA, Osaka, Japan) and allowed to polymerize overnight at 4 °C. After polymerization, the resin block was trimmed with a razor blade and mounted onto a wooden block using adhesive.

Sections of 10 μm thickness were cut with a rotary microtome (HM360; Thermo Fisher Scientific, Waltham, MA, United States), stained with 0.05% aqueous toluidine blue (waldeck GmbH & Co. KG, Germany) and boric acid (Wako) solution, and washed with water. For detailed observation of the root tip shown in Supplementary Figure 1, sections were stained with 0.02% aqueous Safranin O solution. Sections were embedded in Entelan New (Sigma-Aldrich, Darmstadt, Germany). Then, sections were mounted on glass slides and observed with an optical microscopy (DM4500; Leica Microsystems). Fiji/ImageJ software was used for image analysis.

### Quantification of endogenous phytohormones

For the quantification of endogenous phytohormones, approximately 100 mg (fresh weight) of whole roots were collected from plants cultured for two weeks. For C. palustris, low-hardness MS medium solidified with 0.15% gellan gum was used. For *L. arcuata*, Gamborg’s B5 medium solidified with 0.3% gellan gum was used, as B5 medium produces a softer gel than MS medium. Collected root samples were frozen by liquid nitrogen and homogenized using TissueLyser II (QIAGEN, Hilden, Germany). Stable isotope-labeled internal standards were added at the beginning of the extraction procedure. Phytohormones were extracted, semi-purified, and separated into three fractions as previously described (Kojima and Sakakibara. 2012), without MS probe derivatization. ABA and GAs were analyzed by ultra-performance liquid chromatography (UPLC) using an octadecyl-silica (ODS) column (ACQUITY Premier HSS T3 with VanGuard FIT, 1.8 µm, 2.1 × 100 mm; Waters, Milford, MA, USA) coupled to a tandem quadrupole mass spectrometer (ACQUITY UPLC System/Xevo TQ-XS; Waters) equipped with an electrospray ionization (ESI) source. Each fraction was analyzed in separate runs under optimized LC–MS/MS conditions.

### Statistical analysis

All statistical analyses and data visualization were performed using Python (v3.13.2) with the following libraries: pandas for data handling, NumPy for numerical operations, matplotlib.pyplot and seaborn for plotting, and SciPy (scipy.stats) and Pingouin were used for statistical analysis. For comparisons between the terrestrial versus the submerged conditions within C. *palustris*, Welch’s t-test was used. For comparisons among multiple experimental conditions within a species, one-way analysis of variance followed by the Games–Howell post hoc test was applied. For multiple pairwise comparisons of two conditions across different *Callitriche* species, p values were adjusted using the Benjamini–Hochberg false discovery rate (FDR) correction. Error bars in all figures represent the standard error of the mean (SEM). The number of biological replicates (n) is provided in the corresponding figure legends, together with the exact significance thresholds used for each analysis.

## Results

### Phenotypic plasticity of root hair development in *C. palustris*

To observe root phenotype non-invasively, *C. palustris* was grown on a sterilized solid medium under terrestrial and submerged conditions (Fig. 1A). *C. palustris* formed adventitious roots from nodes after cutting and developed few lateral roots on our culture system (Fig 1B). At first glance, roots grown under the two conditions differed drastically in their root hair phenotypes. Based on their growth environments, we hereafter refer to these roots as "terrestrial root" (Ter) and "submerged root" (Sub), respectively. While terrestrial roots had dense long root hairs, submerged root had sparse short root hairs (Fig 1C). Quantifications using a confocal microscope confirmed that all root hair length, density, and hair cell ratio out of all epidermal cells are significantly higher in the terrestrial condition (p < 0.01, Welch’s t-test, Fig. 1D-F).

### The effect of mechanical stimulus and root hair development

Since the physical properties of artificial growth media provide a highly uniform environment that may influence root development through reduced mechanical stimulus, they might not accurately reflect the more heterogeneous conditions of natural soil environments. To evaluate whether the suppression of root hair development observed under laboratory conditions is just caused by the growth on artificial media, we next examined *C. palustris* roots not only under various artificial culture conditions but also on sand under terrestrial and submerged treatments (Fig. 2). First, roots were grown on solid media with different gellan gum concentrations in the terrestrial condition (0.3% and 0.15%), or in a hydroponic system (0% gellan gum), in which shoots remained terrestrial while roots were immersed in liquid medium (Supplementary Fig. 1A). Root hair length and density were reduced on the hydroponic culture condition, whereas they were similar between the 0.15% and standard 0.3% gellan gum media (Games-Howell test, p < 0.05, Fig. 2A, Supplementary Fig. 1B-D). In these conditions, the plants did not show the strong reduction of root hairs comparable to those observed under submerged conditions in the initial experiment.

**Figure 2.**
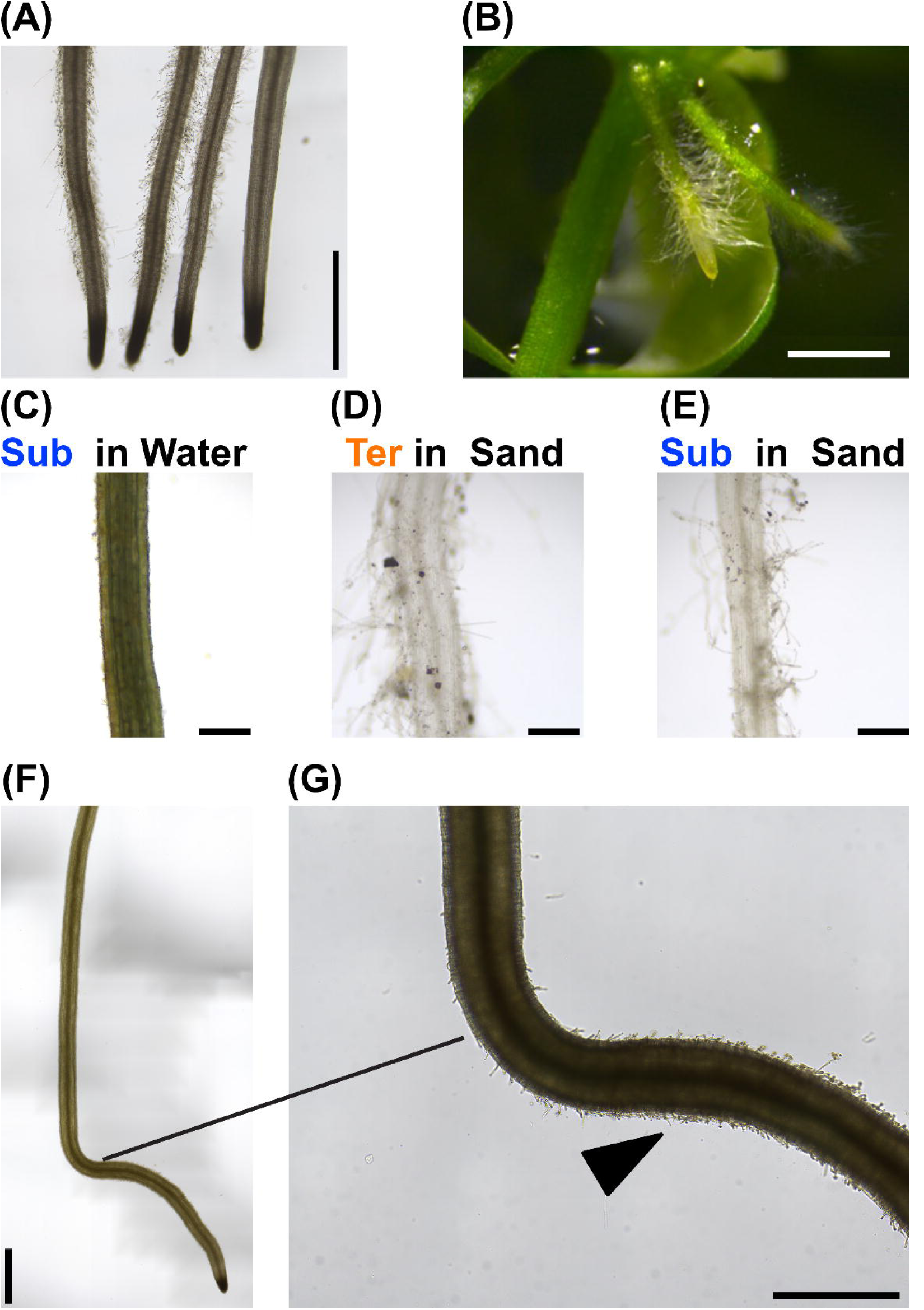
Root hair development under various growth conditions. **(A)** Roots grown in media solidified with various gellan gum concentrations. From left to right in Figure: terrestrial conditions with 0.3%, 0.15%, and 0% gellan gum, respectively, followed by the submerged condition. **(B)** Roots exposed to air above the solid medium. **(C)** Roots developed freely in water in an aquarium tank. **(D)** Terrestrial roots embedded in sand. **(E)** Submerged roots embedded in sand. **(F)** Submerged roots that had grown through the solid medium and reached the bottom surface of the culture box. **(G)** Magnified image of **(F)**. Scale bars: **(A, B, F)** 1 cm; **(C–E)** 200 μm; **(G)** 500 μm.

In the standard terrestrial condition, *C. palustris* also developed roots that emerged in air without contacting solid medium, and such roots had abundant root hairs (Fig. 2B). In contrast, submerged plants in an aquarium tank had roots that grow freely in the water column, which had almost no root hairs (Fig. 2C). In contrast, roots embedded in sand developed root hairs, regardless of whether the shoots were in terrestrial or submerged conditions. (Fig. 2D, E). Finally, roots growing on standard submerged solid medium but mechanically contacting the bottom surface of the culture box exhibited root hair formation at the contact site (Fig. 2F, G).

Across these conditions, root hairs were generally present when shoots remained terrestrial. When shoots were submerged, root hair development was strongly reduced in a solid medium or water. However, root hairs were present even under submerged conditions when roots had physical contact with hard substrates such as the bottom of the culture box or sand particles.

### Phenotypic plasticity of internal root structure in *C. palustris*

In addition to the clear differences in root hair development, submerged roots appeared visibly thicker than terrestrial roots. To determine whether this morphological difference reflects changes in internal anatomy, we made transverse sections from both conditions (Fig. 3A). The root anatomy of *C. palustris* was characterized by a cartwheel-like cortical structure and contained a substantial proportion of intercellular air spaces, forming a well-developed aerenchyma. Also, no obvious positional patterning of root hair formation, as seen in Arabidopsis (Dolan et al. 1993; Salazar-Henao et al. 2016), was detected in the epidermis (Fig. 3A, B). Observation of the longitudinal section around the root tip revealed that cortical layers were sequentially established one by one from the outer to the inner side through successive periclinal cell divisions, a developmental pattern typical in eudicots (Supplementary Fig. 2A, B, Dolan et al. 1993; Heimsch & Seago, 2008). Further examination of transverse serial sections demonstrated that intercellular spaces were generated through cell separation of cortical cells along the circumferential direction, indicating that the aerenchyma formation primary proceeds via a schizogenous process (Supplementary Fig. 2C, D). In addition, some cortical cells exhibited a collapsed morphology, indicative of lysigenous process in the later stage (Supplementary Fig. 2E).

**Figure 3.**
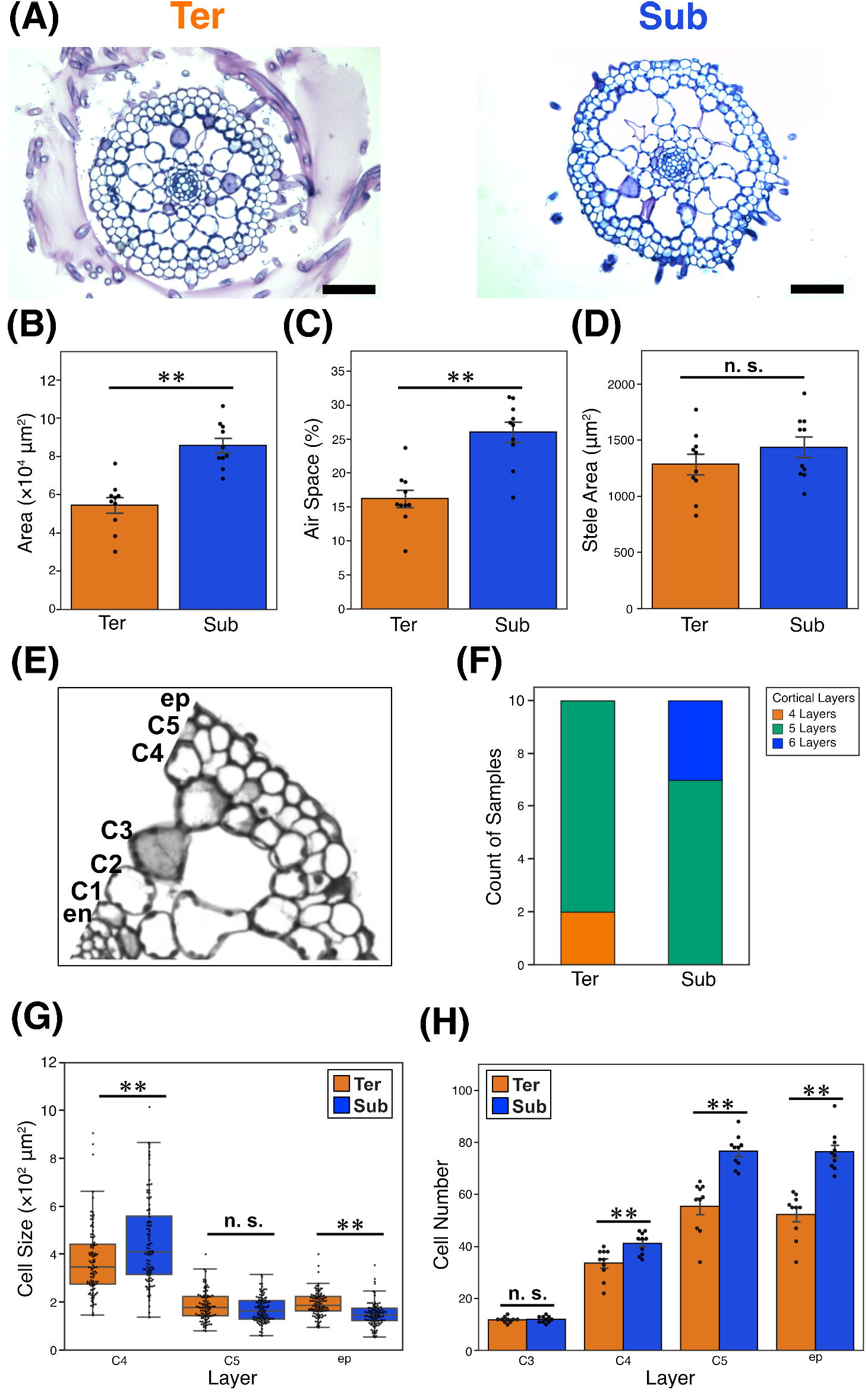
Morphological plasticity of internal root structure in *C. palustris*. **(A)** Cross-sections of roots grown under the two conditions. Scale bars: 100 μm. **(B)** Cross-sectional area. **(C)** Proportion of air space to the total root area. **(D)** Stele area. **(E)** Definitions of cortical layer labels. “en” and “ep” refers endodermis and epidermis, respectively. **(F)** Number of cortical cell layers under the two conditions. n = 10. **(G)** Differences in cell size between the two conditions. n = 100. **(H)** Differences in cell number within each layer under the two conditions. n = 10. Error bars indicate the standard error of the mean (SEM). Asterisks indicate significant differences between conditions (n.s.: not statistically significant; ∗: p < 0.05; ∗∗: p < 0.01; Welch’s t-test).

Submerged roots showed a markedly larger cross-sectional area and a higher proportion of air space than terrestrial roots (p < 0.01, Welch’s t-test, Fig. 3B, C). To determine the underlying cause of this root thickening, we examined the stele area and cortical layer organization (Fig. 3D-F). The stele area did not differ significantly between the two conditions (p > 0.05, Welch’s t-test, Fig. 3D). Cortical layers were typically composed of five layers in both conditions, although submerged roots showed a slight tendency to have more layers (Fig. 3F). Notably, even when samples with the same cortical layer number (five) were compared, significant differences in the total root area or proportion of air space were detected (p < 0.01, Welch’s t-test, Supplementary Fig. 3A-C), indicating that variation in cortical layer number alone does not account for the observed root thickening. Based on the consistent layer organization, we designated the cortical layers outside endodermis as “C1” – “C5” for the further quantitative analysis (Fig. 3E). Because the inner cortical layers contained collapsed cells and air space, analyses were restricted to the outer two cortical layers and the epidermis, where no collapsed cell was observed. Cell sizes in the outer layers showed only small differences between conditions: cells in C4 tended to be slightly larger in submerged roots, whereas C5 showed no significant difference, and epidermal cells were slightly larger in terrestrial roots (p < 0.01, Welch’s t-test, Fig 3G). In contrast, more prominent differences were observed in cell numbers. The number of cells in C3, where aerenchyma is formed, was around 12 cells and almost similar in both conditions (p > 0.05, Welch’s t-test, Fig. 3H). In both environments, the outer layers without aerenchyma (C4–C5 and ep) contained more than twice as many cells as C3, reflecting the higher proliferative activity in outer layers (Fig. 3H). Notably, compared with terrestrial roots, this outer-layer cell number was further increased—by approximately 1.2 to 1.5-fold—in submerged roots (p < 0.01, Welch’s t-test, Fig 3H). Thus, root thickening under the submerged conditions is primarily attributable to this additional increase in cell number rather than cell size or cortical layer number increase. In fact, the root tip was rounder in submerged roots (Supplementary Fig. 3D, E), suggesting enhanced cell proliferation from early stages of root development in the root apical meristem.

Taken together, these morphological data demonstrate pronounced morphological plasticity in root development, which we hereafter term “heterorhizy,” by analogy with heterophylly.

### ABA regulates the plasticity of root hair development

Previous studies have shown that some phytohormones are involved in the regulation of heterophylly (Nakayama et al. 2017; Koga et al. 2024). To test whether phytohormones also contribute to the regulation of heterorhizy, we applied exogenous phytohormones and their inhibitors, focusing on abscisic acid (ABA), gibberellin, and ethylene, which have been implicated in the regulation of heterophylly in *C. palustris* (Koga et al. 2021). Among the hormones tested, ABA had a pronounced effect on root hair development, whereas ethylene, gibberellin, and their inhibitors showed only minor effects (Supplementary Fig. 4). Under the terrestrial condition, the ABA antagonistic inhibitor PANMe (Takeuchi et al. 2018) reduced root hair length and density, whereas exogenous ABA application increased root hair length and density under the submerged condition (Games-Howell test, p < 0.05, Fig. 4A-C). Furthermore, co-treatments demonstrated specificity of ABA action: ABA supplementation rescued the PANMe-induced reduction of root hairs in terrestrial roots, and PANMe abolished ABA-induced root hair formation in submerged roots (Fig. 4B-C, Supplementary Fig. 5). On the other hand, root hair length of ABA treated samples in the submerged condition was shorter than the terrestrial condition (Games-Howell test, p < 0.05, Fig. 4B).

**Figure 4.**
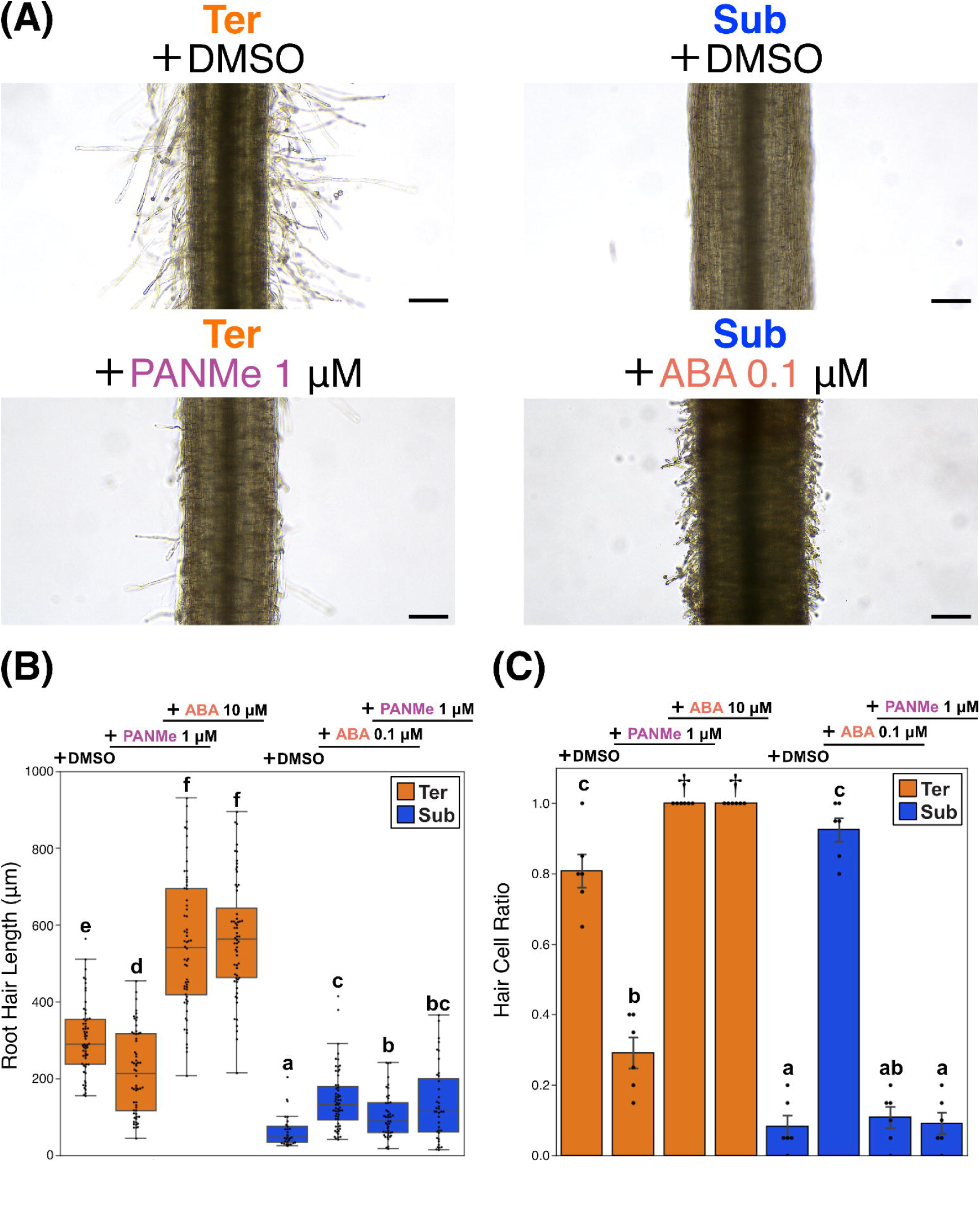
ABA promotes root hair development. **(A)** Effect of ABA and PANMe (ABA antagonistic inhibitor) treatments on root hair development. Scale bar: 100 μm. **(B)** Root hair length under treatments. n = 29−60. **(C)** Hair cell ratio under treatments. n = 6. Error bars indicate the standard error of the mean (SEM). Different letters indicate statistically significant differences among groups according to the Games–Howell test (p < 0.05). †: All measured values were equal to 1.0; therefore, statistical comparison was not applicable.

We next quantified endogenous phytohormone levels. To prevent interference with fresh weight measurements caused by adhesion of the culture medium to root hairs, we used a solid medium containing a low concentration of gellan gum (0.15% (w/v)) used in the previous experiment. Endogenous ABA content tended to be higher under the terrestrial condition, whereas the difference was not statistically significant (Welch’s t-test, p > 0.05, Supplementary Fig. 5C).

### GA regulates cell proliferation and root thickening

We also examined transverse sections of roots treated with the three phytohormones. Root area was largely unchanged by ABA, ethylene, or their inhibitors (Supplementary Fig. 6), whereas treatment with GA₃ or the GA biosynthesis inhibitor uniconazole P (Uni) led to clear changes in root area (Games-Howell test, p < 0.05, Fig. 5A-B). GA₃ treatment reduced root area under both terrestrial and submerged conditions, while uniconazole P increased it in both conditions. A higher concentration of GA₃ reversed the effect of uniconazole P (Fig. 5B-C, Supplementary Fig. 7). These changes in root diameter were accompanied by corresponding alterations in cell number (Games-Howell test, p < 0.05, Fig. 5C). GA₃ treatment reduced the proportion of air space under the submerged condition, whereas uniconazole P did not cause a clear increase in air space formation in either environment (Games-Howell test, p > 0.05, Supplementary Fig. 7B). We also attempted to quantify endogenous GAs. However, neither GA₁ nor GA₄ was quantified in our measurements, as their levels were below the quantification limit of the method (1 fmol for both GA1 and GA4).

**Figure 5.**
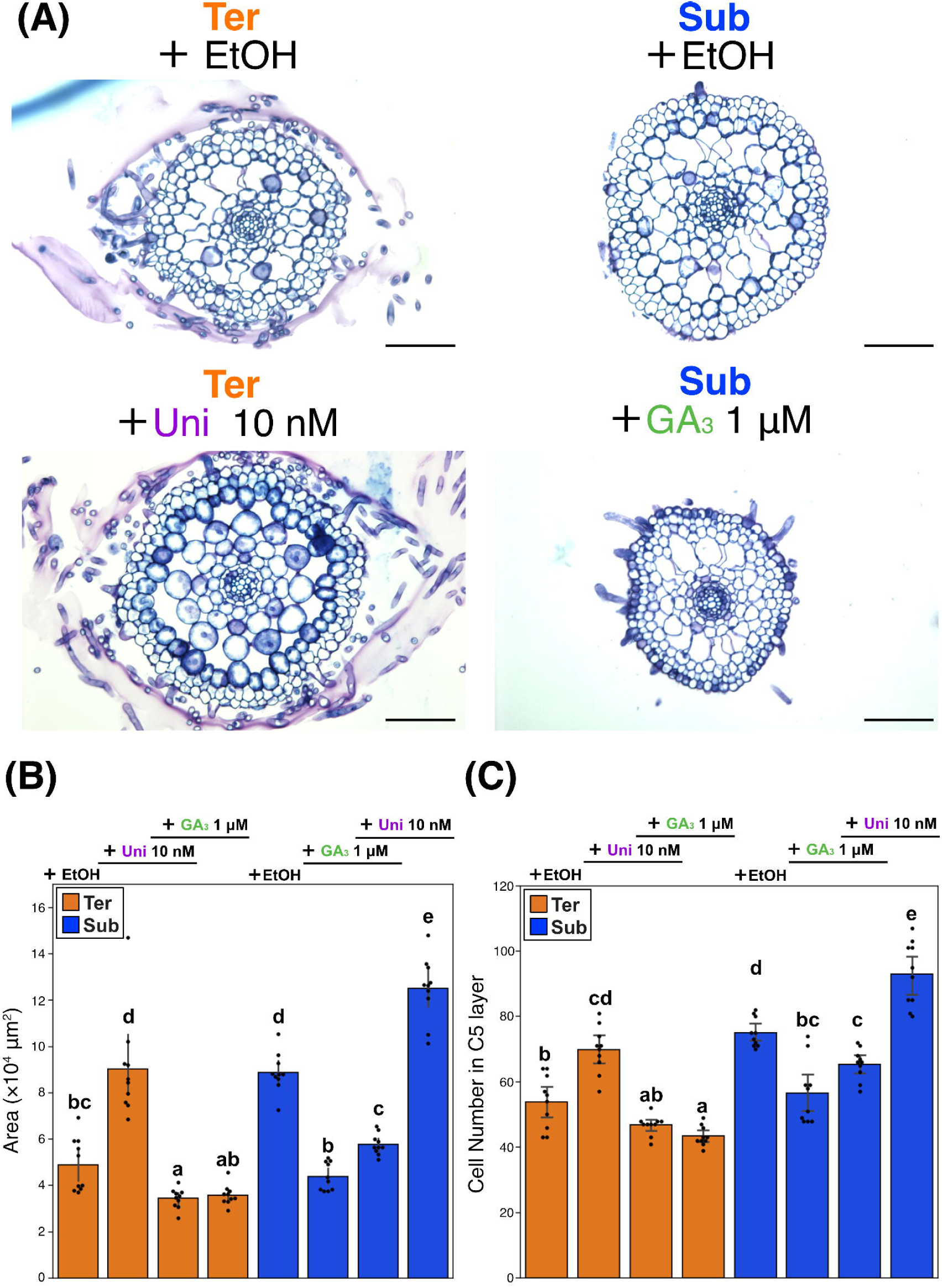
Gibberellin negatively regulates root thickening via cell division. **(A)** Effect of GA_3_ and Uniconazole P treatments on root anatomy. Scale bar: 100 μm. **(B)** Root cross-sectional area under treatments. **(C)** Epidermal cell number under treatments. n = 10. Error bars indicate the standard error of the mean (SEM). Different letters indicate statistically significant differences among groups according to the Games–Howell test (p < 0.05).

### Assessment of the morphological plasticity in other *Callitriche species*

To reveal the evolutionary process of heterorhizy in *C. palustris*, we examined whether relative species of *Callitriche palustris* have similar plasticity in response to submergence. Four other *Callitriche* species inhabiting in Japan (*C. japonica, C. stagnalis, C. terrestris, and C. deflexa*) were used in this study (Fig 6A). *C. japonica* is a terrestrial species occupying a basally diverged phylogenetic position; *C. stagnalis* is an amphibious species; whereas *C. terrestris* and *C. deflexa* are secondarily terrestrial species that evolved from aquatic ancestors (Fig 6A, Ito et al. 2017, Koga et al. 2024). Reduction of root hairs and substantial root anatomical changes in the submerged condition were observed only in the amphibious species *C. stagnalis*, in which cell number in the C5 layer increased by approximately 20 cells (Welch’s t-test with FDR correction, q < 0.05, Fig. 6A, B, Supplementary Fig. 8). In contrast, *C. japonica*, *C. terrestris*, and *C. deflexa* exhibited milder yet statistically significant changes in root thickness, air space proportion, and cell number, with the C5 cortical layer increasing by only about five cells. (Welch’s t-test with FDR correction, q < 0.05, Fig. 6A, C, Supplementary Fig. 9). Whereas collapsed cortical cells were detected in the other species, they were not observed in *C. japonica* (Supplementary Fig. 9A).

**Figure 6.**
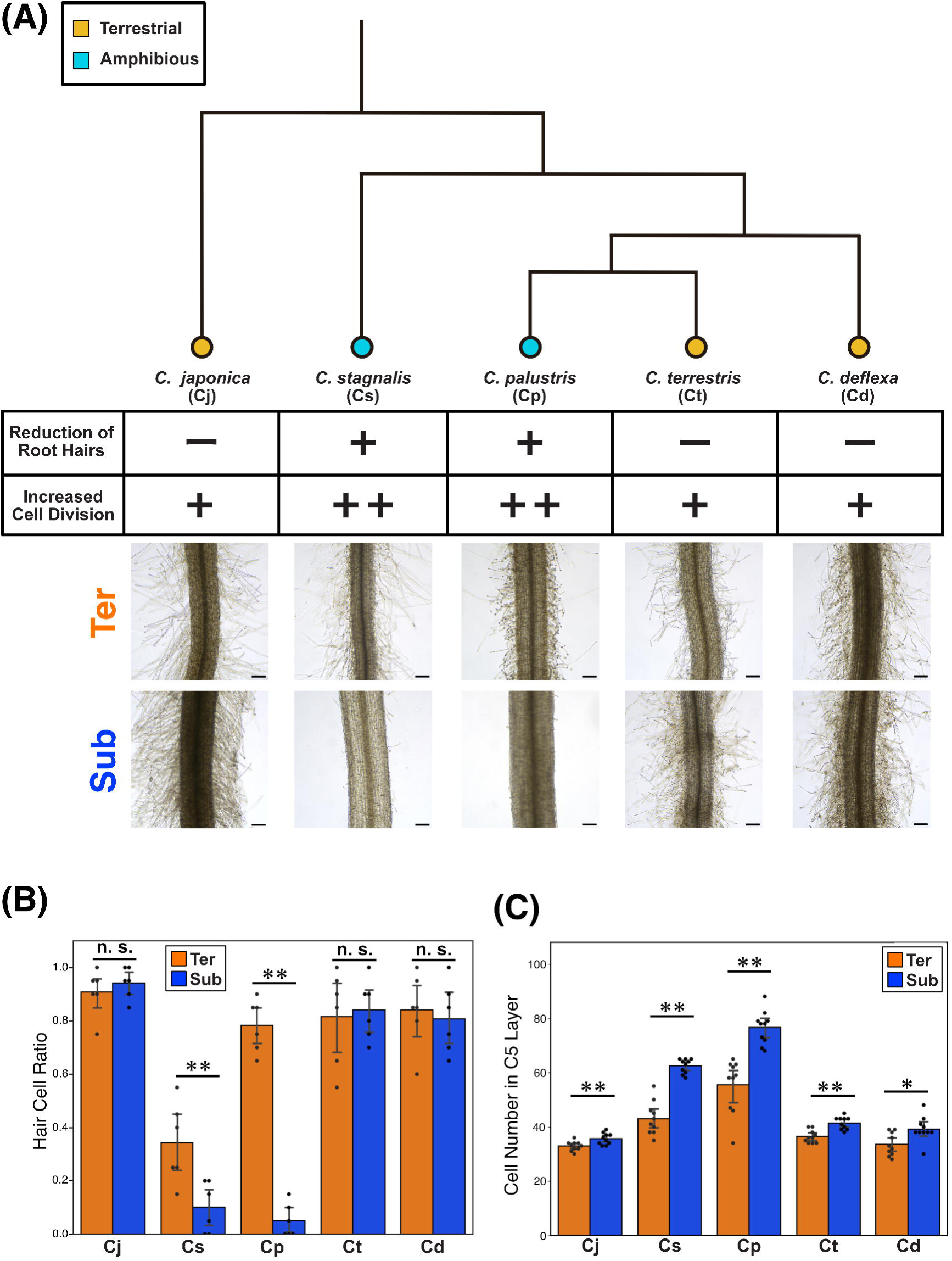
Root morphological plasticity in relative *Callitriche* species. **(A)** Simplified phylogenetic tree of genus *Callitriche* and the presence pattern of submergence-induced morphological plasticity in five species. ++: Drastic morphological plasticity; +: Mild morphological plasticity; -: No morphological plasticity. The phylogenetic tree was constructed based on Ito et al. (2017) and Koga et al. (2024). **(B)** Difference of hair cell ratio under two conditions. n = 6. **(C)** Difference of epidermal cell number under two conditions. n = 10. Error bars indicate the standard error of the mean (SEM). Asterisks indicate significant differences between conditions based on Welch’s t-test followed by FDR (Benjamini–Hochberg) correction (n.s.: not statistically significant; ∗: p < 0.05; ∗∗: p < 0.01.).

### Heterorhizy in a distinct amphibious plant *Ludwigia arcuata*

To examine whether morphological plasticity like *C. palustris* is also seen in distinct amphibious plants, we investigated *Ludwigia arcuata* (Onagraceae, Myrates). Despite being phylogenetically distant from Callitriche, this amphibious species exhibits pronounced heterophylly, producing leaf morphologies similar to those of *C. palustris*. (Fig. 7A, Kuwabara et al. 2001, 2003; The Angiosperm Phylogeny Group, 2016). Under sterilized solid-medium conditions, *L. arcuata* showed clear root hair plasticity similar to that of *C. palustris*, developing abundant root hairs under the terrestrial conditions while producing no root hairs under the submerged conditions (Fig. 7B). Furthermore, PANMe treatment under the terrestrial conditions suppressed root hair development, whereas ABA treatment under the submerged conditions promoted root hair development, in a manner comparable to that observed in *C. palustris* (Fig. 7B, C, Supplementary Fig. 10A). The endogenous ABA content in the roots of *L. arcuata* also tended to be higher under terrestrial conditions, although the difference was not statistically significant (Welch’s t-test, p > 0.05, Supplementary Fig. 10B).

**Figure 7.**
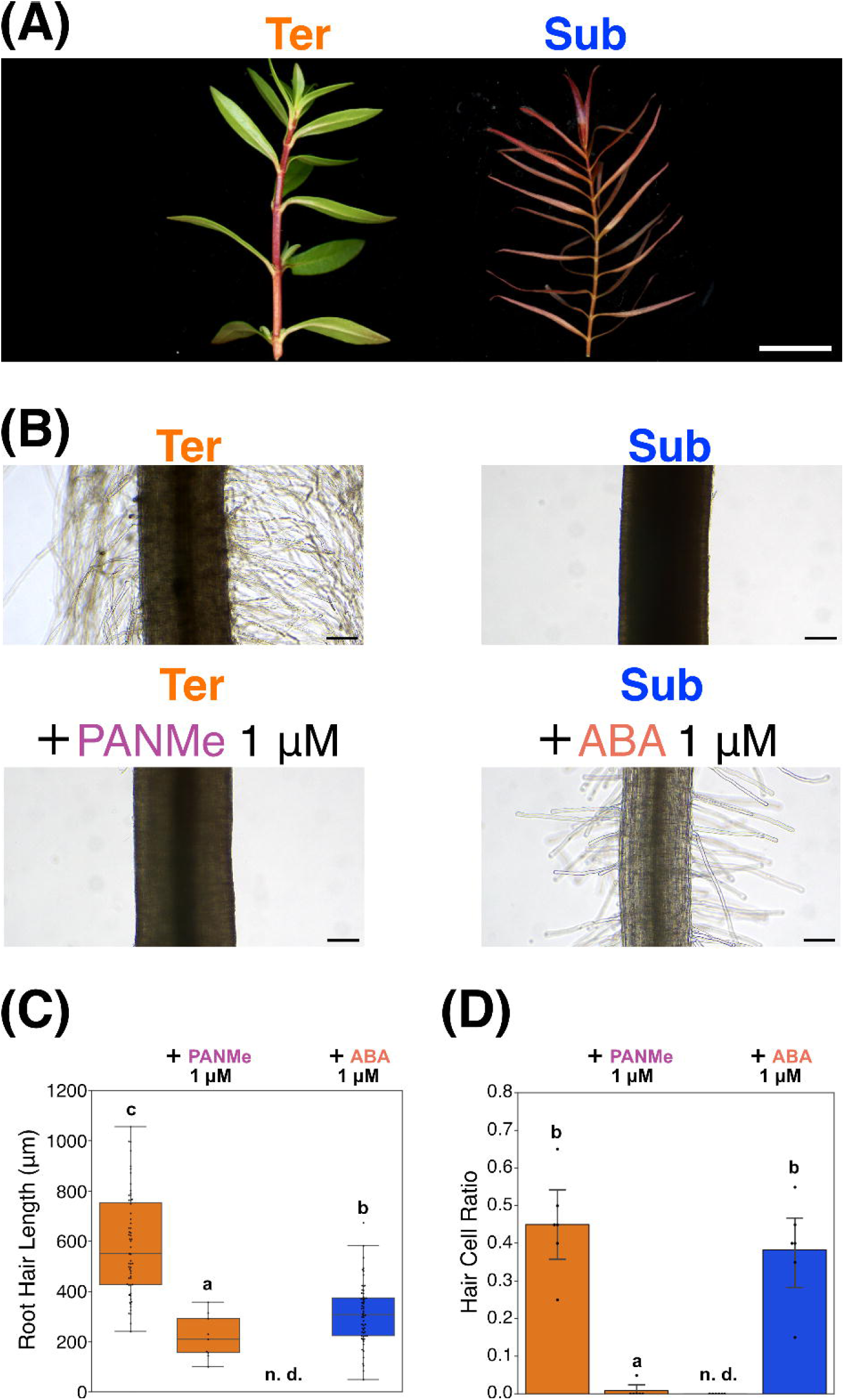
Heterorhizy in *Ludwigia arcuata*. **(A)** Heterophylly in *L. arcuata*. **(B)** The plasticity of root hair and the effect on phytohormone in *L. arcuata*. **(C)** Root hair length under different conditions. n = 9-60. **(D)** Hair cell ratio under different conditions. n = 6. Different letters indicate statistically significant differences among groups according to the Games–Howell test (p < 0.05). Scale bars: **(A)**: 1 cm; **(B)**: 100 μm.

### Root traits similar to the submerged root of *C. palustris* in diverse aquatic plants

To assess how widespread root morphologies similar to those of submerged root in *C. palustris* are among angiosperms, we further examined the morphology of adventitious roots in several aquatic plant species and conducted a phylogenetically broad survey to identify species exhibiting similar root traits (Supplementary Tables 2). We identified submerged-root phenotypes with few or no root hairs in the roots of several aquatic species across angiosperms, such as *Rotala hippuris* (Haloragaceae, Myrates), *Myriophyllum spicatum* (Lythraceae, Saxifragales), and *Egeria densa* (Hydrocharitaceae, Alismatales) (Fig. 8A, Supplementary Fig. 11). While several species developed few root hairs in water, they developed root hairs after penetration to soil, similar to *C. palustris* and previously documented species (Supplementary Fig. 11, Shannon, 1953). Moreover, several aquatic species such as *Hydrocotyle verticillate* (Araliaceae, Apiales), *Rorippa aquatica* (Brassicaceae, Brassicales), and *Elatine triandra* (Elatinaceae, Malpighiales) developed cartwheel-like schizogenous aerenchyma (Fig. 8B). Although morphological plasticity in response to submergence was not explicitly examined in the species, the number of cells in the cortical layer outside the air space was approximately twice that of the air space-harboring layer (Fig. 8B), whose radial organization was similar to that of *C. palustris*.

**Figure 8.**
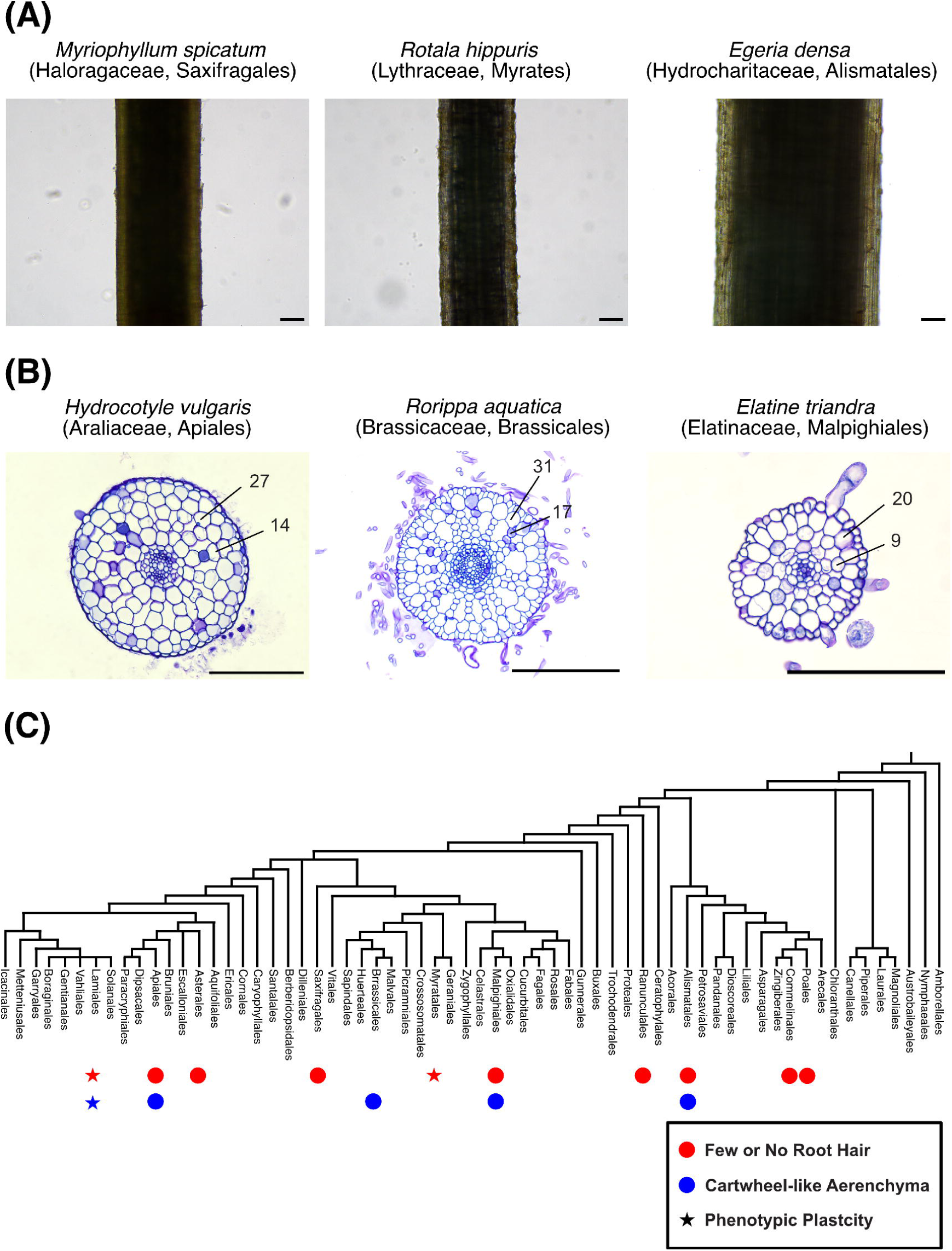
Evolutionary generality of suppression of root hair development and cartwheel-like aerenchyma formation in various aquatic plants. **(A)** Roots with few root hairs in three aquatic plants. **(B)** Cartwheel-like aerenchyma formation in three aquatic plants. Cell numbers are indicated for the outermost aerenchyma-harboring layer and the innermost non-aerenchyma-harboring layer. **(C)** Phylogenetic distribution of sparse root hair phenotype and cartwheel-like aerenchyma formation in aquatic angiosperms. Phylogenetic tree was constructed based on APG IV system (Angiosperm Phylogeny Group, 2016). Scale bars: **(A)**: 100 μm; **(B)**: 50 μm.

**Figure 9.**
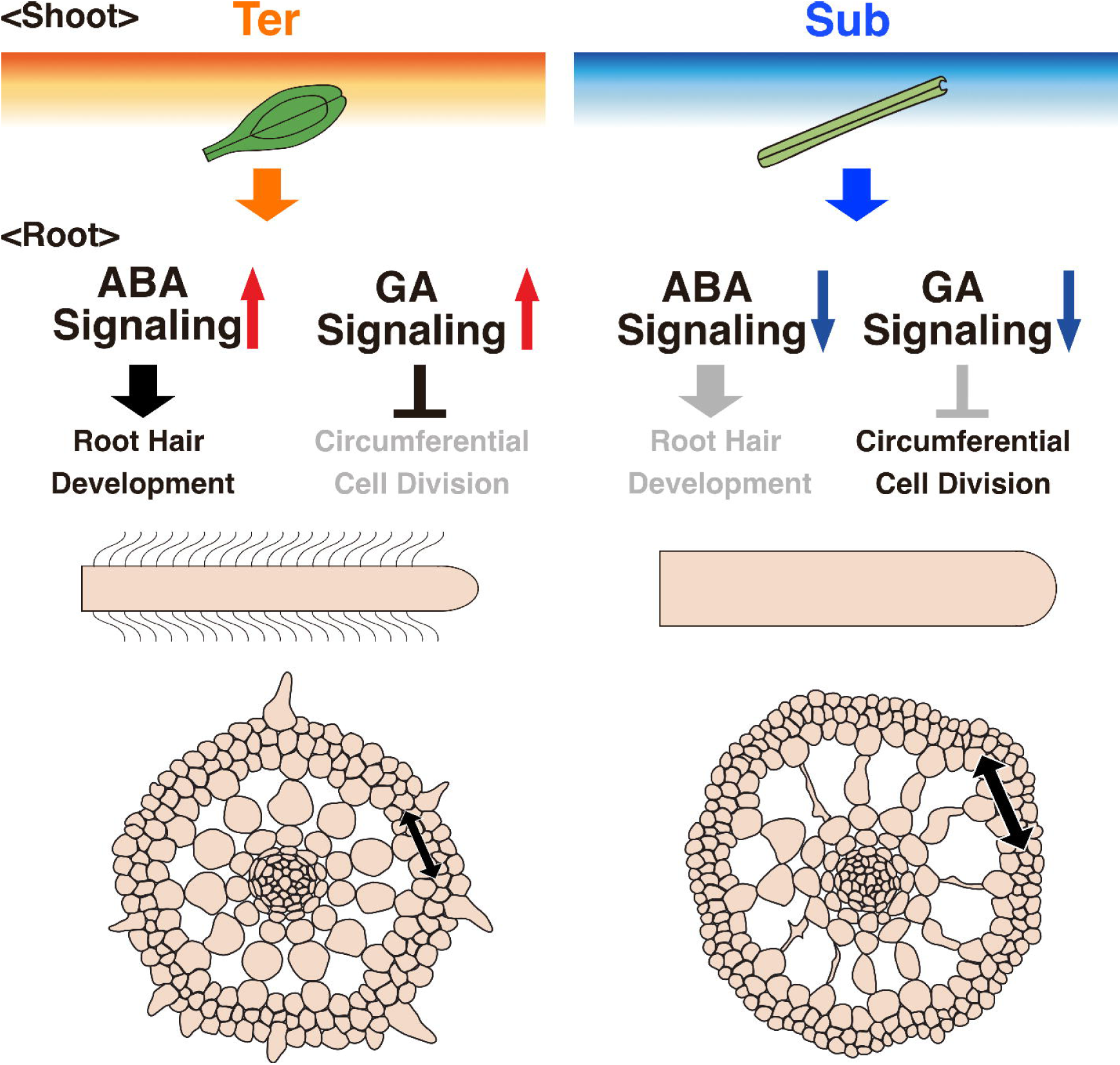
Schematic figure of *C. palustris* heterorhizy.

To place these observations in a broader phylogenetic context, we mapped currently confirmed presence of sparse root hair and cartwheel-type aerenchyma formation in aquatic plants across the angiosperm phylogeny, incorporating both our own observations (Supplementary Table 2) and previously reported cases compiled from the literature (Supplementary Table 3; Shannon, 1953; Clayton & Bagyaraj, 1984; Ware et al., 2023; Yin et al., 2025). Both sparse or no root hair formation and cartwheel-like aerenchyma were observed across multiple, phylogenetically distant aquatic lineages, indicating that these submerged root traits seen in *C. palustris* are shared in diverse aquatic plants.

## Discussion

### “Heterorhizy” in the amphibious plant *C. palustris*

We found that *Callitriche palustris* exhibits remarkable morphological plasticity in root hair development and root anatomy. Inspired by the concept of heterophylly in amphibious plants, we termed this plasticity in root morphology “heterorhizy”. Historically, the term “heterorhizy” has been used to describe the phenomenon in which a single plant produces morphologically distinct types of roots, such as in rice lateral roots, often in association with environmental or developmental cues (Tschirch, 1905; Kumazawa, 1979; Sutton, 1980; Montagnoli et al. 2014; Watanabe et al. 2020). In this study, we extend the meaning of heterorhizy to encompass submergence-induced root morphological plasticity in amphibious plants. Although amphibious plants have been studied in the context of heterophylly, heterorhizy in amphibious plants has long been overlooked. We propose heterorhizy as a fundamental concept of morphological plasticity in amphibious plants, comparable in importance to heterophylly.

### Speculated nature and physiological advantage of the plasticity of root hair development

Previous studies reported that several aquatic plants do not produce root hairs (Cormack, 1937; Shannon, 1953; Yin et al. 2025), but the dramatic change in root hair development of amphibious plants has been scarcely investigated. Our results including experiments in solid medium and other conditions indicate that shoot submergence itself is a key trigger for the suppression of root hair development, because the physical and chemical conditions surrounding the root were similar between terrestrial and submerged conditions in solid medium. On the other hand, mechanical stimulus to roots also promoted root hair formation even in the submerged condition. This result is consistent with earlier observations showing that some aquatic plants develop root hairs only when their roots contact soil, but not when submerged freely in water (Cormack, 1937; Shannon, 1953). Taken together, our solid-medium-based experimental system appears to mimic conditions in which roots are submerged but not in contact with the substrate, as in roots suspended in the water column.

We speculate that the suppression of root hair development under the submerged condition may reduce the unnecessary metabolic cost associated with producing root hairs, because water can be absorbed through the shoot in submerged plants (Chambers et al. 1989; Madsen & Cedergreen, 2002; Kuntz et al. 2014). In contrast, the induction of root hair development by mechanical contact suggests that root hairs may play functional roles such as physical anchorage to soil. Further investigation will be required to clarify the physiological and ecological advantage of the suppression of root hair development.

### Mechanisms of the plasticity in schizogenous aerenchyma formation in *C. palustris*

Waterlogging-induced aerenchyma formation has been extensively studied in Poaceae species such as rice and maize, where it typically occurs through a lysigenous process by programmed cell death (Yamauchi et al. 2013; Takahashi et al. 2014). While lysigenous cavities were also observed in *C. palustris,* our analyses revealed a distinct developmental process: enhanced cell proliferation in the outer cortical layers contributes to the expansion of schizogenous aerenchyma. Although cortical layers C3–C5 originate from a common lineage, the outer layers lacking air spaces (C4, C5) consistently contain more than twice as many cells as C3, the layer containing air spaces, indicating preferential circumferential proliferation in the outer cortex after layer establishment. The resulting imbalance in circumferential expansion likely increases tensile forces on the inner layers, producing cartwheel-like schizogenous aerenchyma. A comparable morphology has been described in a wetland Brassicaceae species *Cardamine amara* (Kudoh et al. 2025), whereas submergence-induced plasticity in this structure has not been reported.

Our observations suggest that *C. palustris* employs the same fundamental developmental strategy of layer-specific cell proliferation under both terrestrial and submerged conditions, whereas the proliferation is further enhanced under the submerged condition. The similar number of cells in C3 under terrestrial and submerged roots indicates that the number of the original cortical cell lineage is comparable between two conditions. However, submergence increases the frequency of circumferential cell divisions in the C4 and C5 layers after cortical layer establishment, leading to pronounced radial thickening of the root and further expansion of schizogenous aerenchyma. These findings propose another form of plasticity in aerenchyma formation, which is distinct from the well-known lysigenous mechanism.

### Phytohormone-mediated regulation of heterorhizy

ABA showed a strong positive effect on the development of root hair not only in *C. palustris*, but also in *L. arcuata*. ABA is well known for its roles in drought stress responses and has also been implicated in promoting aerial leaf phenotypes in amphibious plants (Anderson, 1978; Goliber & Feldman, 1989; Kuwabara et al. 2003; Li et al. 2017; Kim et al. 2018; Koga et al. 2021; Sakamoto et al. 2024). Thus, ABA appears to pleiotropically contribute to terrestrial traits in both heterophylly and heterorhizy of *C. palustris*. Previous studies have shown that ABA suppresses root hair elongation in *Arabidopsis thaliana* (Rymen et al. 2017), whereas it promotes root hair elongation in *Oryza sativa* (Wang et al. 2017). Together with previous reports, our results indicate that a common ABA input can lead to distinct root hair phenotypes among species, underscoring diversity in the downstream execution of ABA signaling during root development.

*C. palustris* increases overall root thickness by increasing the number of cortical and epidermal cells and the proliferation was negatively regulated by GA. Previous studies revealed that GA represses root thickening by repressing cambial cell division in radish root (Meng et al. 2024), or producing slender cells in *Medicago truncatula* and *Lactuca sativa* (Tanimoto, 1987; Fonouni-Farde et al. 2019). GA also represses the layer-increasing periclinal divisions in the root cortex of *A. thaliana* and *M. truncatula* (Gong et al. 2016; Fonouni-Farde et al. 2019). While GA typically affects root thickness through changes in cell size or layer-increasing division in other species, GA modulates circumferential proliferation within a given cortical layer in *C. palustris*. The difference indicates that, even when the input of GA signal and the output of root thickness are similar, the intervening cellular processes differ among species.

Our results indicate that ABA and GA have distinct effects on heterorhizy, with ABA affecting root hair development and GA affecting root cell division. This functional separation suggests that ABA- and GA-dependent responses are deployed in different downstream cascades of submergence. On the other hand, our measurements did not detect a significant difference of ABA contents and did not detect endogenous gibberellin in roots. This might suggest that these phytohormones are localized within a restricted region of the root tip, or that their signaling outputs are modulated without substantial changes in hormone production. A similar situation has been reported for heterophylly in *C. palustris*, where GA signaling contributes to leaf form change despite endogenous GA not being detected in shoots (Koga et al., 2021). Also, ABA treatment under the submerged condition did not recover root hair length as the terrestrial condition, and Uniconazole P treatment did not increase the proportion of air space. These suggest that ABA- and GA-dependent pathways alone are insufficient, and that additional regulatory components in the downstream of submergence is required to completely coordinate heterorhizy.

### Evolution of heterorhizy

The phylogenetic distribution of root morphological plasticity within the genus *Callitriche* indicates that the drastic form of heterorhizy is not uniformly present across the genus but is restricted to amphibious species. The largely unchanged root phenotype of terrestrial species in response to submergence suggests that the plasticity seen in amphibious species is not merely stress-induced developmental impairment. These observations raise the possibility that heterorhizy is a derived trait which evolved within the genus *Callitriche*.

Our results indicate that *Ludwigia arcuata* also exhibits submergence-induced heterorhizy, characterized by root hair plasticity similar to that of *C. palustris*. Notably, in both species, root hair development is regulated by similar ABA response, highlighting the similar regulatory mechanism in these two species. Although the evolutionary origin of heterorhizy of *Ludwigia* remains unresolved, the large phylogenetic distance between *Ludwigia* and *Callitriche* suggests that the observed similarity is more consistent with convergent evolution than with shared ancestry.

In addition to the plasticity in *L. arcuata*, this study and previous studies also observed suppression of root hair development in aquatic plants. Although submerged species such as *Egeria densa* cannot be evaluated for the plasticity between terrestrial and submerged conditions, they similarly lack root hairs in submerged roots. This phylogenetically widespread absence of root hairs suggests that suppression of root hair development under the submerged condition is an important adaptive trait across aquatic plants. Moreover, many of these species formed root hairs after penetration to soil, indicating that they have not completely lost the root hair developmental machinery, unlike duckweeds (Ware et al. 2023). Instead, as in submerged roots of *C. palustris*, root hair development in these species is likely suppressed until roots reach to soil. Also, cartwheel-like aerenchyma structure was observed in diverse aquatic plants outside the genus *Callitriche*. The shared morphological feature suggests that a similar developmental program to submerged root development in *C. palustris* has been repeatedly recruited in various aquatic plants. Together, these observations suggest that root features highlighted in submerged roots of *C. palustris* are widespread among diverse aquatic plants.

Importantly, since *C. palustris* have both terrestrial-type root morphology and submerged-type root morphology within a single species, it serves as a powerful model as a transitional state between terrestrial and aquatic root developmental programs. By elucidating the regulatory mechanisms underlying heterorhizy in *C. palustris*, it becomes possible to gain insight into how the widespread aquatic root traits are developmentally regulated. Therefore, heterorhizy in *C. palustris* will be an important framework to understand how root developmental programs are modified during adaptation to aquatic environments.

## Acknowledgements

We are grateful to Asami Ota for technical assistance with plant cultivation. We thank the WINGS-LST program and Life Sciences Core Facility at the University of Tokyo for providing access to confocal microscopy facilities. We also thank Professor Seisuke Kimura for supplying plant material of *Rorippa aquatica.* We acknowledge the use of ChatGPT (OpenAI) for assistance with English language editing and improvement of manuscript clarity.

## Author contributions

T. S., Y. D., H. K., and H. T. designed the research. T. S. and Y. D. carried out the initial observations. T. S. conducted all experiments and analyzed the data except measurement of phytohormone amount. T. S. and H. K. identified the plant species. M. K., Y. T. and H. S. conducted the measurement of phytohormone amount. J. T. and Y. T. performed the synthesis of the ABA antagonist PANMe. All authors contributed to writing the manuscript.

## References

Abiko, T., Kotula, L., Shiono, K., Malik, A. I., Colmer, T. D., & Nakazono, M. (2012). Enhanced formation of aerenchyma and induction of a barrier to radial oxygen loss in adventitious roots of *Zea nicaraguensis* contribute to its waterlogging tolerance as compared with maize (*Zea* mays ssp. Mays). Plant, Cell & Environment, 35(9), 1618‒1630. 10.1111/j.1365-3040.2012.02513.x

Anderson, L. W. J. (1978). Abscisic acid induces formation of floating leaves in the heterophyllous aquatic angiosperm *Potamogeton nodosus*. Science, 201(4361), 1135‒1138. 10.1126/science.201.4361.1135

Arber, A. (1920). Water plants, a study of aquatic angiosperms. Cambridge University Press.

Bailey-Serres, J., & Voesenek, L. A. C. J. (2008). Flooding stress: acclimations and genetic diversity. Annual Review of Plant Biology, 59(1), 313‒339. 10.1146/annurev.arplant.59.032607.092752

Binder, B. M. (2020). Ethylene signaling in plants. Journal of Biological Chemistry, 295(22), 7710‒7725. 10.1074/jbc.REV120.010854

Chambers, P. A., Prepas, E. E., Bothwell, M. L., & Hamilton, H. R. (1989). Roots versus shoots in nutrient uptake by aquatic macrophytes in flowing waters. Canadian Journal of Fisheries and Aquatic Sciences, 46(3), 435‒439. 10.1139/f89-058

Clayton, J. S., & Bagyaraj, D. J. (1984). Vesicular-arbuscular mycorrhizas in submerged aquatic plants of New Zealand. Aquatic Botany, 19(3‒4), 251‒262. 10.1016/0304-3770(84)90043-3

Colmer, T. D. (2003). Long-distance transport of gases in plants: a perspective on internal aeration and radial oxygen loss from roots. Plant, Cell & Environment, 26(1), 17‒36. 10.1046/j.1365-3040.2003.00846.x

Colmer, T. D., Winkel, A., Kotula, L., Armstrong, W., Revsbech, N. P., & Pedersen, O. (2020). Root O_2_ consumption, CO_2_ production and tissue concentration profiles in chickpea, as influenced by environmental hypoxia. New Phytologist, 226(2), 373‒384. 10.1111/nph.16368

Cook, C. D. K. (1999). The number and kinds of embryo-bearing plants which have become aquatic: a survey. Perspectives in Plant Ecology, Evolution and Systematics, 2(1), 79‒102. 10.1078/1433-8319-00066

Cormack, R. G. H. (1937). The development of root hairs by *Elodea canadensi*s. New Phytologist, 36(1), 19‒25. 10.1111/j.1469-8137.1937.tb06900.x

Delwiche, C. F., & Cooper, E. D. (2015). The evolutionary origin of a terrestrial flora. Current Biology, 25(19), R899‒910. 10.1016/j.cub.2015.08.029

Dolan, L., Janmaat, K., Willemsen, V., Linstead, P., Poethig, S., Roberts, K., & Scheres, B. (1993). Cellular organisation of the *Arabidopsis thaliana* root. Development, 119(1), 71‒84. 10.1242/dev.119.1.71

Doll, Y., Koga, H., & Tsukaya, H. (2021). *Callitriche* as a potential model system for evolutionary studies on the dorsiventral distribution of stomata. Plant Signaling & Behavior, 16(11), 1978201. 10.1080/15592324.2021.1978201

Fonouni-Farde, C., Miassod, A., Laffont, C., Morin, H., Bendahmane, A., Diet, A., & Frugier, F. (2019). Gibberellins negatively regulate the development of *Medicago truncatula* root system. Scientific Reports, 9(1), 2335. 10.1038/s41598-019-38876-1

Fukao, T., Barrera-Figueroa, B. E., Juntawong, P., & Peña-Castro, J. M. (2019). Submergence and waterlogging stress in plants: a review highlighting research opportunities and understudied aspects. Frontiers in Plant Science, 10, 340. 10.3389/fpls.2019.00340

Goliber, T. E., & Feldman, L. J. (1989). Osmotic stress, endogenous abscisic acid and the control of leaf morphology in *Hippuris vulgaris* L. *Plant*, Cell & Environment, 12(2), 163‒171. 10.1111/j.1365-3040.1989.tb01929.x

Gong, X., Flores-Vergara, M. A., Hong, J. H., Chu, H., Lim, J., Franks, R. G., Liu, Z., & Xu, J. (2016). SEUSS integrates gibberellin signaling with transcriptional inputs from the SHR-SCR-SCL3 module to regulate middle cortex formation in the Arabidopsis root. Plant Physiology, 170(3), 1675‒1683. 10.1104/pp.15.01501

Guo, L., Yin, L., Sun, C., Zhao, K., Zhao, H., Bai, S.-N., Li, Y., & Wu, W. (2025). Gradual genomic streamlining and convergent adaptation during terrestrial-to-aquatic transitions in angiosperms. Current Biology, 35 (19), 4595‒4605. 10.1016/j.cub.2025.08.001

Heimsch, C., & Seago, J. L. (2008). Organization of the root apical meristem in angiosperms. American Journal of Botany, 95(1), 1‒21. 10.3732/ajb.95.1.1

Horiguchi, G., Nemoto, K., Yokoyama, T., & Hirotsu, N. (2019). Photosynthetic acclimation of terrestrial and submerged leaves in the amphibious plant *Hygrophila difformis*. AoB PLANTS, 11(2). 10.1093/aobpla/plz009

Ito, Y., Tanaka, N., Barfod, A. S., Kaul, R. B., Muasya, A. M., Garcia-Murillo, P., De Vere, N., Duyfjes, B. E. E., & Albach, D. C. (2017). From terrestrial to aquatic habitats and back again: Molecular insights into the evolution and phylogeny of *Callitriche* (Plantaginaceae). Botanical Journal of the Linnean Society, 184(1), 46‒58. 10.1093/botlinnean/box012

Jung, J., Lee, S. C., & Choi, H.-K. (2008). Anatomical patterns of aerenchyma in aquatic and wetland plants. Journal of Plant Biology, 51(6), 428‒439. 10.1007/BF03036065

Kim, J., Joo, Y., Kyung, J., Jeon, M., Park, J. Y., Lee, H. G., Chung, D. S., Lee, E., & Lee, I. (2018). A molecular basis behind heterophylly in an amphibious plant, *Ranunculus trichophyllus*. PLOS Genetics, 14(2), e1007208. 10.1371/journal.pgen.1007208

Koga, H., Doll, Y., Hashimoto, K., Toyooka, K., & Tsukaya, H. (2020). Dimorphic leaf development of the aquatic plant *Callitriche palustris* L. through differential cell division and expansion. Frontiers in Plant Science, 11, 269. 10.3389/fpls.2020.00269

Koga, H., Ikematsu, S., & Kimura, S. (2024). Diving into the Water: Amphibious Plants as a Model for Investigating Plant Adaptations to Aquatic Environments. Annual Review of Plant Biology, 75(1), 579‒604. 10.1146/annurev-arplant-062923-024919

Koga, H., Kojima, M., Takebayashi, Y., Sakakibara, H., & Tsukaya, H. (2021). Identification of the unique molecular framework of heterophylly in the amphibious plant *Callitriche palustris* L. The Plant Cell, 33(10), 3272‒3292. 10.1093/plcell/koab192

Kojima M, Sakakibara H. (2012). Highly sensitive high-throughput profiling of six phytohormones using MS-probe modification and liquid chromatography‒tandem mass spectrometry. In: Normanly, J. (ed.) High-Throughput Phenotyping in Plants. Methods in Molecular Biology, 918, 151‒164. 10.1007/978-1-61779-995-2_11

Kudoh, H., Sakane, M., Marhold, K., Shimizu-Inatsugi, R., Shimizu, K. K., & Fukaki, H. (2025). Cartwheel aerenchyma in *Cardamine amara* as a model of schizogenous tissue formation in plants. iScience, 28(8), 113106. 10.1016/j.isci.2025.113106

Kumazawa, M. (1979). Plant organography. Shokabo.

Kuntz, K., Heidbüchel, P., & Hussner, A. (2014). Effects of water nutrients on regeneration capacity of submerged aquatic plant fragments. Annales de Limnologie - International Journal of Limnology, 50(2), 155‒162. 10.1051/limn/2014008

Kurihara, D., Mizuta, Y., Sato, Y., & Higashiyama, T. (2015). ClearSee: A rapid optical clearing reagent for whole-plant fluorescence imaging. Development, 142 (23), 4168‒4179. 10.1242/dev.127613

Kuwabara, A., Ikegami, K., Koshiba, T., & Nagata, T. (2003). Effects of ethylene and abscisic acid upon heterophylly in *Ludwigia arcuata* (Onagraceae). Planta, 217(6), 880‒887. 10.1007/s00425-003-1062-z

Kuwabara, A., Tsukaya, H., & Nagata, T. (2001). Identification of factors that cause heterophylly in *Ludwigia arcuata* Walt. (Onagraceae). Plant Biology, 3(1), 98‒105. 10.1055/s-2001-11748

Li, G., Hu, S., Hou, H., & Kimura, S. (2019). Heterophylly: phenotypic plasticity of leaf shape in aquatic and amphibious plants. Plants, 8(10), 420. 10.3390/plants8100420

Li, G., Hu, S., Yang, J., Schultz, E. A., Clarke, K., & Hou, H. (2017). Water-Wisteria as an ideal plant to study heterophylly in higher aquatic plants. Plant Cell Reports, 36(8), 1225‒1236. 10.1007/s00299-017-2148-6

Lughadha, E. N., Govaerts, R., Belyaeva, I., Black, N., Lindon, H., Allkin, R., Magill, R. E., & Nicolson, N. (2016). Counting counts: Revised estimates of numbers of accepted species of flowering plants, seed plants, vascular plants and land plants with a review of other recent estimates. Phytotaxa, 272(1), 82. 10.11646/phytotaxa.272.1.5

Madsen, T. V., & Cedergreen, N. (2002). Sources of nutrients to rooted submerged macrophytes growing in a nutrient-rich stream. Freshwater Biology, 47(2), 283‒291. 10.1046/j.1365-2427.2002.00802.x

Meng, G., Yong, M., Zhang, Z., Zhang, Y., Wang, Y., Xiong, A., & Su, X. (2024). Exogenous gibberellin suppressed taproot secondary thickening by inhibiting the formation and maintenance of vascular cambium in radish (*Raphanus sativus* L.). Frontiers in Plant Science, 15, 1395999. 10.3389/fpls.2024.1395999

Mommer, L., & Visser, E. J. W. (2005). Underwater photosynthesis in flooded terrestrial plants: A matter of leaf plasticity. Annals of Botany, 96(4), 581‒589. 10.1093/aob/mci212

Montagnoli, A., Terzaghi, M., Scippa, G. S., & Chiatante, D. (2014). Heterorhizy can lead to underestimation of fine-root production when using mesh-based techniques. Acta Oecologica, 59, 84‒90. 10.1016/j.actao.2014.06.004

Nakayama, H., Nakayama, N., Seiki, S., Kojima, M., Sakakibara, H., Sinha, N., & Kimura, S. (2014). Regulation of the KNOX-GA gene module induces heterophyllic alteration in north American lake cress. The Plant Cell, 26(12), 4733‒4748. 10.1105/tpc.114.130229

Nakayama, H., Sinha, N. R., & Kimura, S. (2017). How do plants and phytohormones accomplish heterophylly, leaf phenotypic plasticity, in response to environmental cues. Frontiers in Plant Science, 8, 1717. 10.3389/fpls.2017.01717

Nielsen, S. L. (1993). A comparison of aerial and submerged photosynthesis in some Danish amphibious plants. Aquatic Botany, 45(1), 27‒40. 10.1016/0304-3770(93)90050-7

Preibisch, S., Saalfeld, S., & Tomancak, P. (2009). Globally optimal stitching of tiled 3D microscopic image acquisitions. Bioinformatics, 25(11), 1463‒1465. 10.1093/bioinformatics/btp184

Rademacher, W. (2000). Growth retardants: effects on gibberellin biosynthesis and other metabolic pathways. Annual Review of Plant Physiology and Plant Molecular Biology, 51(1), 501‒531. 10.1146/annurev.arplant.51.1.501

Rymen, B., Kawamura, A., Schäfer, S., Breuer, C., Iwase, A., Shibata, M., Ikeda, M., Mitsuda, N., Koncz, C., Ohme-Takagi, M., Matsui, M., & Sugimoto, K. (2017). ABA suppresses root hair growth via the OBP4 transcriptional regulator. Plant Physiology, 173(3), 1750‒1762. 10.1104/pp.16.01945

Sakamoto, Tomoaki, Ikematsu, S., Nakayama, H., Mandáková, T., Gohari, G., Sakamoto, Takuya, Li, G., Hou, H., Matsunaga, S., Lysak, M. A., & Kimura, S. (2024). A chromosome-level genome assembly for the amphibious plant *Rorippa aquatica* reveals its allotetraploid origin and mechanisms of heterophylly upon submergence. Communications Biology, 7(1), 431. 10.1038/s42003-024-06088-7

Salazar-Henao, J. E., Vélez-Bermúdez, I. C., & Schmidt, W. (2016). The regulation and plasticity of root hair patterning and morphogenesis. Development, 143(11), 1848‒1858. 10.1242/dev.132845

Sanderson, M. J., Thorne, J. L., Wikström, N., & Bremer, K. (2004). Molecular evidence on plant divergence times. American Journal of Botany, 91(10), 1656‒1665. 10.3732/ajb.91.10.1656

Schenk, H. (1889). Ueber das Aerenchym, ein dem Kork homologes Gewebe bei Sumpflanzen. Jahrbucher Fur Wissenschaftliche Botanik, Jahrbucher Fur Wissenschaftliche Botanik, 20, 526‒574.

Schindelin, J., Arganda-Carreras, I., Frise, E., Kaynig, V., Longair, M., Pietzsch, T., Preibisch, S., Rueden, C., Saalfeld, S., Schmid, B., Tinevez, J.-Y., White, D. J., Hartenstein, V., Eliceiri, K., Tomancak, P., & Cardona, A. (2012). Fiji: An open-source platform for biological-image analysis. Nature Methods, 9(7), 676‒682. 10.1038/nmeth.2019

Sculthorpe, C. D. (1967). The Biology of Aquatic Vascular Plants. Edward Arnold.

Seago, J. L., Marsh, L. C., Stevens, K. J., Soukup, A., Votrubová, O., & Enstone, D. E. (2005). A Re-examination of the root cortex in wetland flowering plants with respect to aerenchyma. Annals of Botany, 96(4), 565‒579. 10.1093/aob/mci211

Shannon, E. L. (1953). The production of root hairs by aquatic plants. American Midland Naturalist, 50(2), 474. 10.2307/2422106

Smirnoff, N., & Crawford, R. M. M. (1983). Variation in the structure and response to flooding of root aerenchyma in some wetland plants. Annals of Botany, 51(2), 237‒249. 10.1093/oxfordjournals.aob.a086462

Sutton, R. F. (1980). Root system morphogenesis. New Zealand Journal of Forestry Science, 10, 264‒292.

Takahashi, H., Yamauchi, T., Colmer, T. D., & Nakazono, M. (2014). Aerenchyma formation in plants. In: Van Dongen JT, Licausi F, eds. Low-Oxygen Stress in Plants. Springer, 247‒265. 10.1007/978-3-7091-1254-0_13

Takeuchi, J., Mimura, N., Okamoto, M., Yajima, S., Sue, M., Akiyama, T., Monda, K., Iba, K., Ohnishi, T., & Todoroki, Y. (2018). Structure-based chemical design of abscisic acid antagonists that block PYL‒PP2C receptor interactions. ACS Chemical Biology, 13(5), 1313‒1321. 10.1021/acschembio.8b00105

Tanimoto, E. (1987). Gibberellin-dependent root elongation in *Lactuca sativa*: recovery from growth retardant-suppressed elongation with thickening by low concentration of GA_3_. Plant and Cell Physiology, 28(6), 963‒973. 10.1093/oxfordjournals.pcp.a077399

The Angiosperm Phylogeny Group. (2016). An update of the Angiosperm Phylogeny Group classification for the orders and families of flowering plants: APG IV. Botanical Journal of the Linnean Society, 181(1), 1‒20. 10.1111/boj.12385

Tschirch, A. (1905). Über die Heterorhizie bei Dikotylen. Flora oder Allgemeine Botanische Zeitung, 94, 68‒78. 10.1016/S0367-1615(17)31598-7

Wang, T., Li, C., Wu, Z., Jia, Y., Wang, H., Sun, S., Mao, C., & Wang, X. (2017). Abscisic acid regulates auxin homeostasis in rice root tips to promote root hair elongation. Frontiers in Plant Science, 8, 1121. 10.3389/fpls.2017.01121

Ware, A., Jones, D. H., Flis, P., Chrysanthou, E., Smith, K. E., Kümpers, B. M. C., Yant, L., Atkinson, J. A., Wells, D. M., Bhosale, R., & Bishopp, A. (2023). Loss of ancestral function in duckweed roots is accompanied by progressive anatomical reduction and a re-distribution of nutrient transporters. Current Biology, 33(9), 1795–1802.e4. 10.1016/j.cub.2023.03.025

Watanabe, Y., Kabuki, T., Kakehashi, T., Kano-Nakata, M., Mitsuya, S., & Yamauchi, A. (2020). Morphological and histological differences among three types of component roots and their differential contribution to water uptake in the rice root system. Plant Production Science, 23(2), 191‒201. 10.1080/1343943X.2020.1730701

Wells, C. L., & Pigliucci, M. (2000). Adaptive phenotypic plasticity: The case of heterophylly in aquatic plants. Perspectives in Plant Ecology, Evolution and Systematics, 3(1), 1‒18. 10.1078/1433-8319-00001

Yamauchi, T., Colmer, T. D., Pedersen, O., & Nakazono, M. (2018). Regulation of root traits for internal aeration and tolerance to soil waterlogging-flooding stress. Plant Physiology, 176(2), 1118‒1130. 10.1104/pp.17.01157

Yamauchi, T., Shimamura, S., Nakazono, M., & Mochizuki, T. (2013). Aerenchyma formation in crop species: A review. *Field Crops Research*, Crop Resilience, 152, 8‒16. 10.1016/j.fcr.2012.12.008

Yamauchi, T., Tanaka, A., Mori, H., Takamure, I., Kato, K., & Nakazono, M. (2016). Ethylene-dependent aerenchyma formation in adventitious roots is regulated differently in rice and maize. Plant, Cell & Environment, 39(10), 2145‒2157. 10.1111/pce.12766

Yamauchi, T., Tanaka, A., Tsutsumi, N., Inukai, Y., & Nakazono, M. (2020). A role for auxin in ethylene-dependent inducible aerenchyma formation in rice roots. Plants, 9(5), 610. 10.3390/plants9050610

Yin, J., Zhu, T., Li, X., Wang, F., & Xu, G. (2025). Phytoremediation of microplastics by water hyacinth. Environmental Science and Ecotechnology, 24, 100540. 10.1016/j.ese.2025.100540

