## Supplemental Figures and Tables for "The submergence-induced drastic morphological plasticity of root in the amphibious plant *Callitriche palustris*"

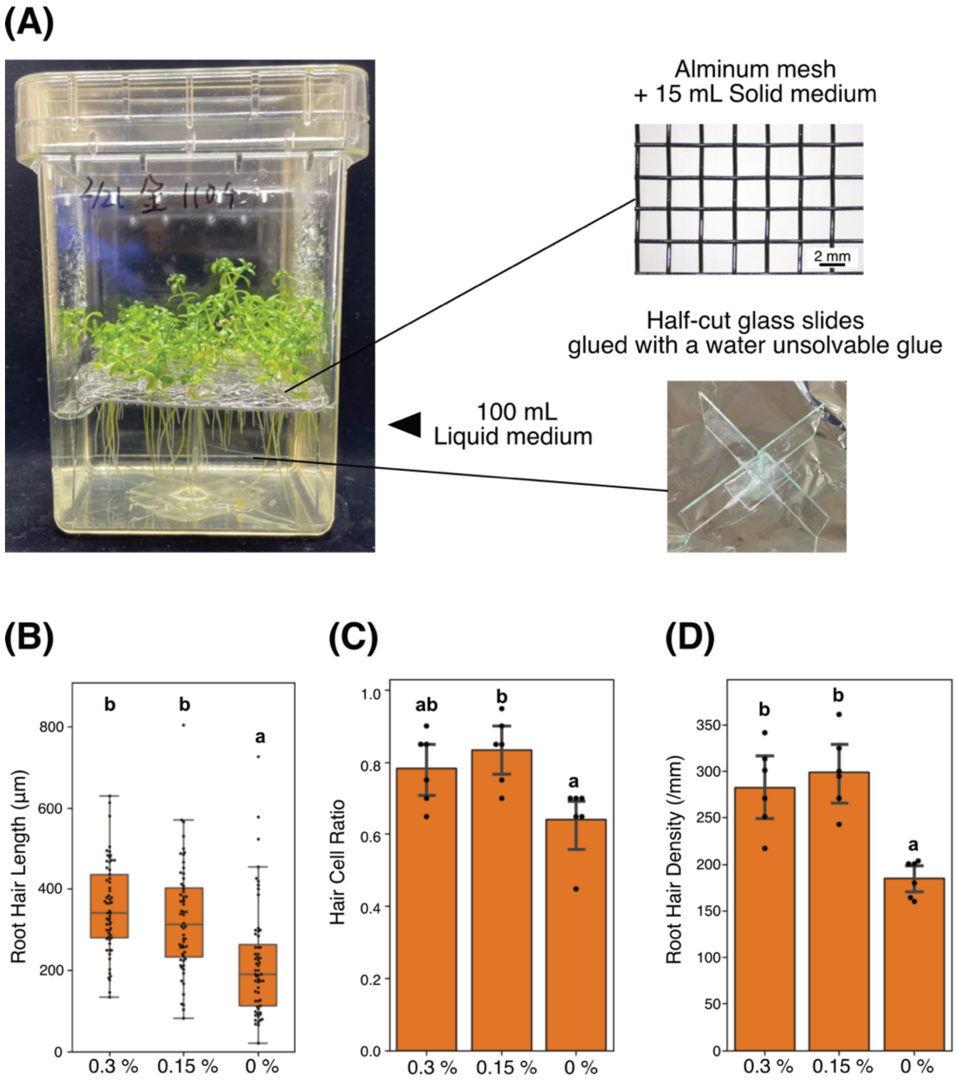


**Supplementary Figure 1.** Scheme of hydroponic culture and quantitative data. **(A)** Scheme of hydroponic culture of *C. palustris.* **(B–D)** Quantitative analyses of root hair traits under there gellan gum concentrations. **(B)** Root hair length. n = 60. **(C)** Proportion of root hair cells among all epidermal cells. n = 6. **(D)** Root hair density. n = 6. Error bars indicate the standard error of the mean (SEM). Different letters indicate statistically significant differences among treatment groups according to the Games–Howell test ($p < 0.05$).

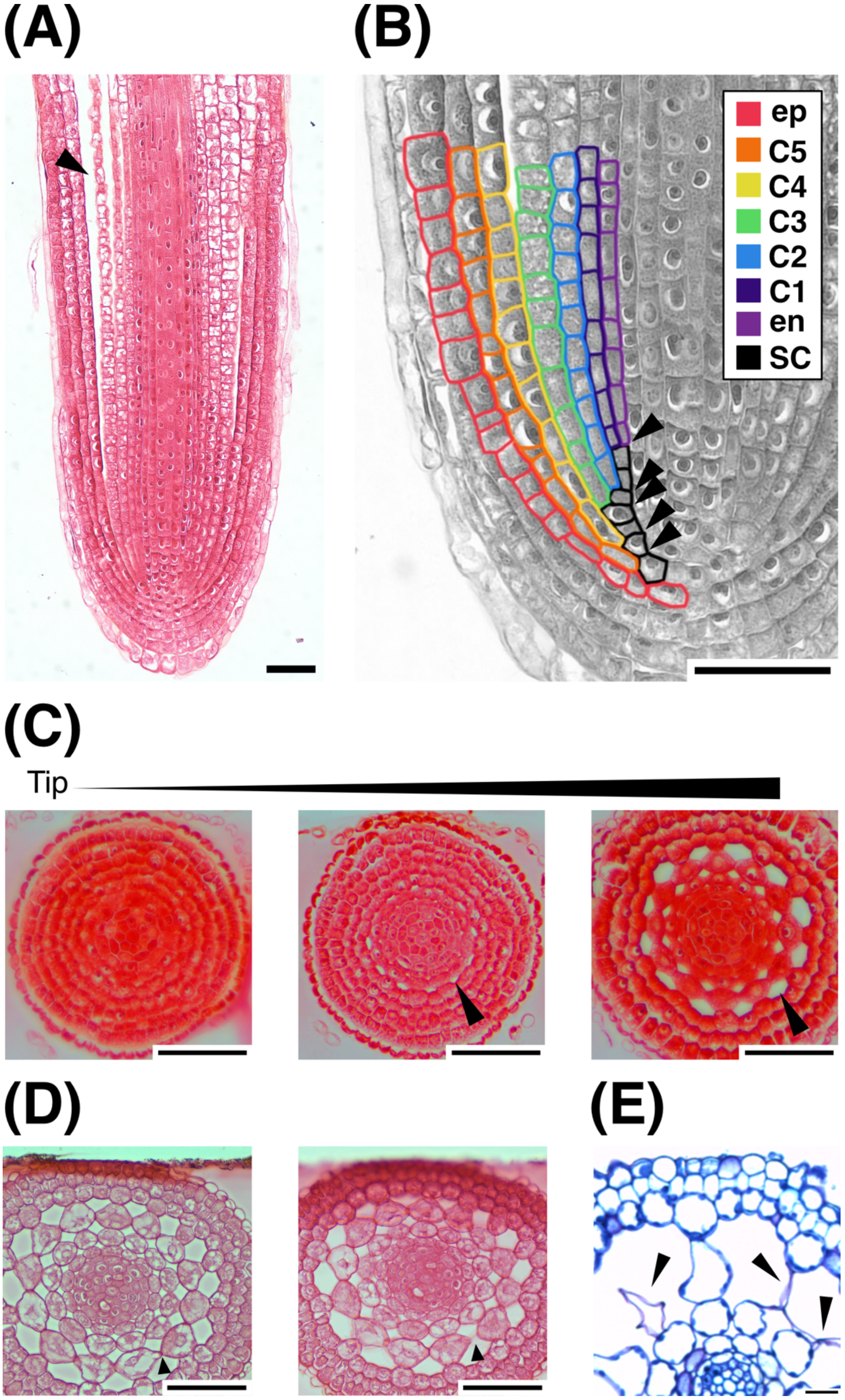


**Supplementary Figure 2.** Aerenchyma formation in *C. palustris*. **(A)** Longitudinal section of a root tip under the submerged condition. Arrowheads indicate air space in aerenchyma. **(B)** Progressive increase in cortical layers via periclinal divisions.

Arrowheads indicates the point of periclinal cell divisions. en, endodermis; C1–C5, cortical layers; ep, epidermis; SC, stem cell. **(C)** Representative sections from a serial section set showing air space formation during early root development. Arrowheads indicate air space. **(D)** Pair of serial transverse sections at 10 μm intervals, showing circumferential cell separation. Arrowheads indicate cell separation. **(E)** Collapsed cortical cells produced by lysigenous process under the submerged condition. Arrowheads indicate collapsed cortical cells. Scale bars: **(A, B)** 10 μm, **(C–E)** 20 μm.

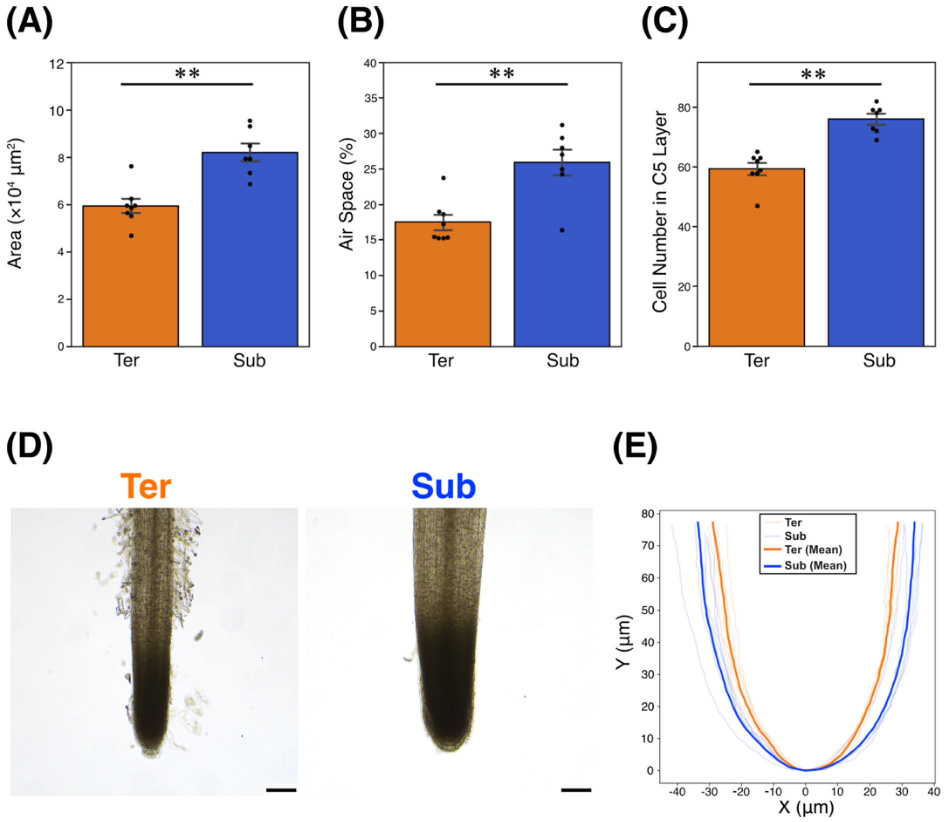


**Supplementary Figure 3.** Comparative analysis of cortical anatomy and root tip outline in terrestrial and submerged roots. **(A–C)** Comparison of terrestrial and submerged root samples with five cortical layers. **(A)** Root cross-sectional area. **(B)** Proportion of air space. **(C)** Epidermal cell number. n = 7. Error bars indicate the standard error of the mean (SEM). Asterisks indicate statistically significant differences between conditions (**: $p < 0.01$; Welch's t-test). **(D)** Root tip morphology under terrestrial and submerged conditions. **(E)** Outlines of root tips derived from six samples per condition. Scale bars: 50 µm.

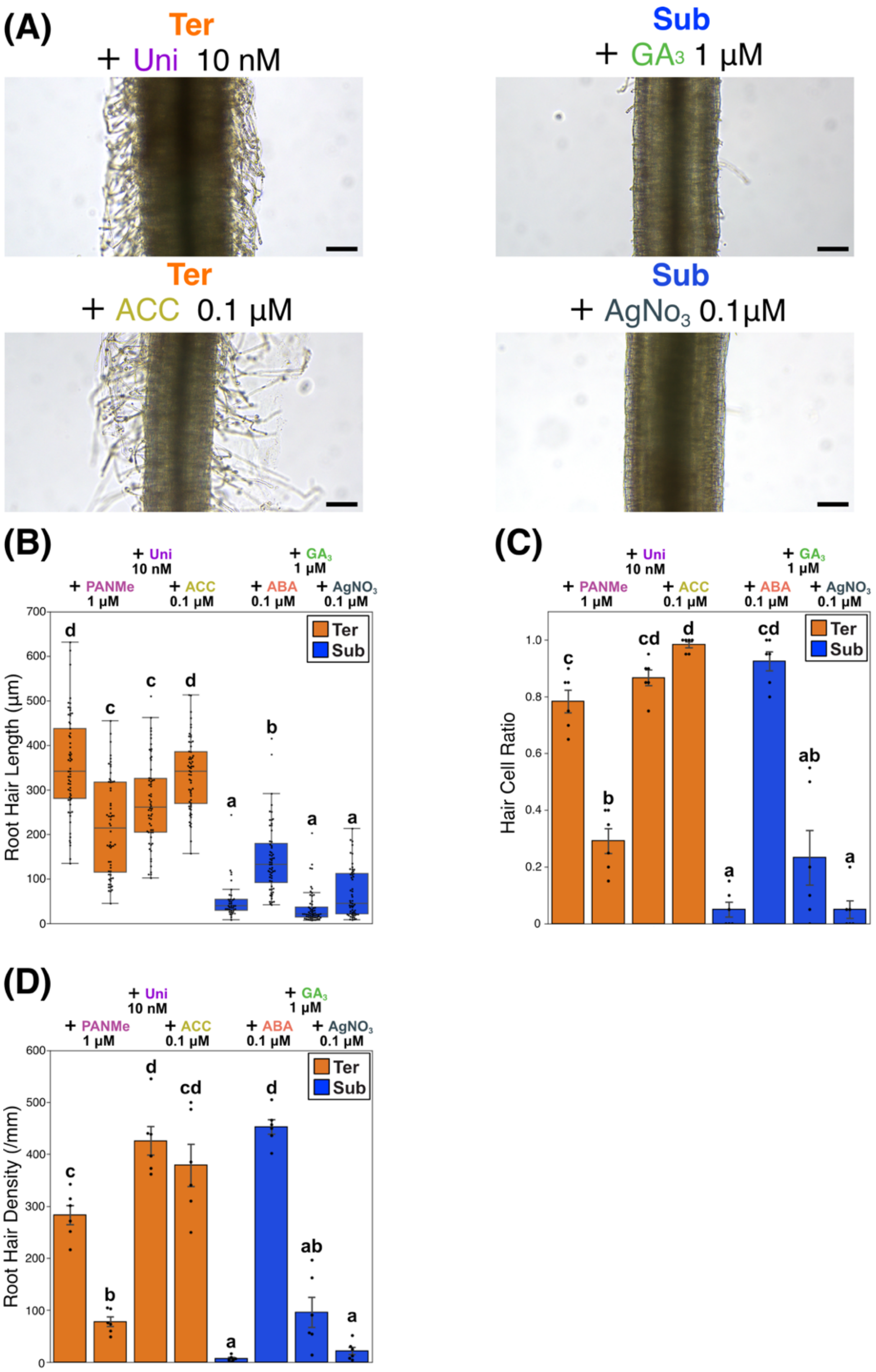


**Supplementary Figure 4.** Effect of phytohormones on root hair development. **(A)** Effects of ethylene, GA, and their inhibitors on root hair development. Scale bar: 100 µm. **(B–D)** Quantitative analyses of root hair traits. **(B)** Root hair length (n = 38–60). **(C)** Hair cell ratio (n = 6). **(D)** Root hair density (n = 6). Error bars indicate the standard error of the mean (SEM). Different letters indicate statistically significant differences among groups according to the Games–Howell test ($p < 0.05$).

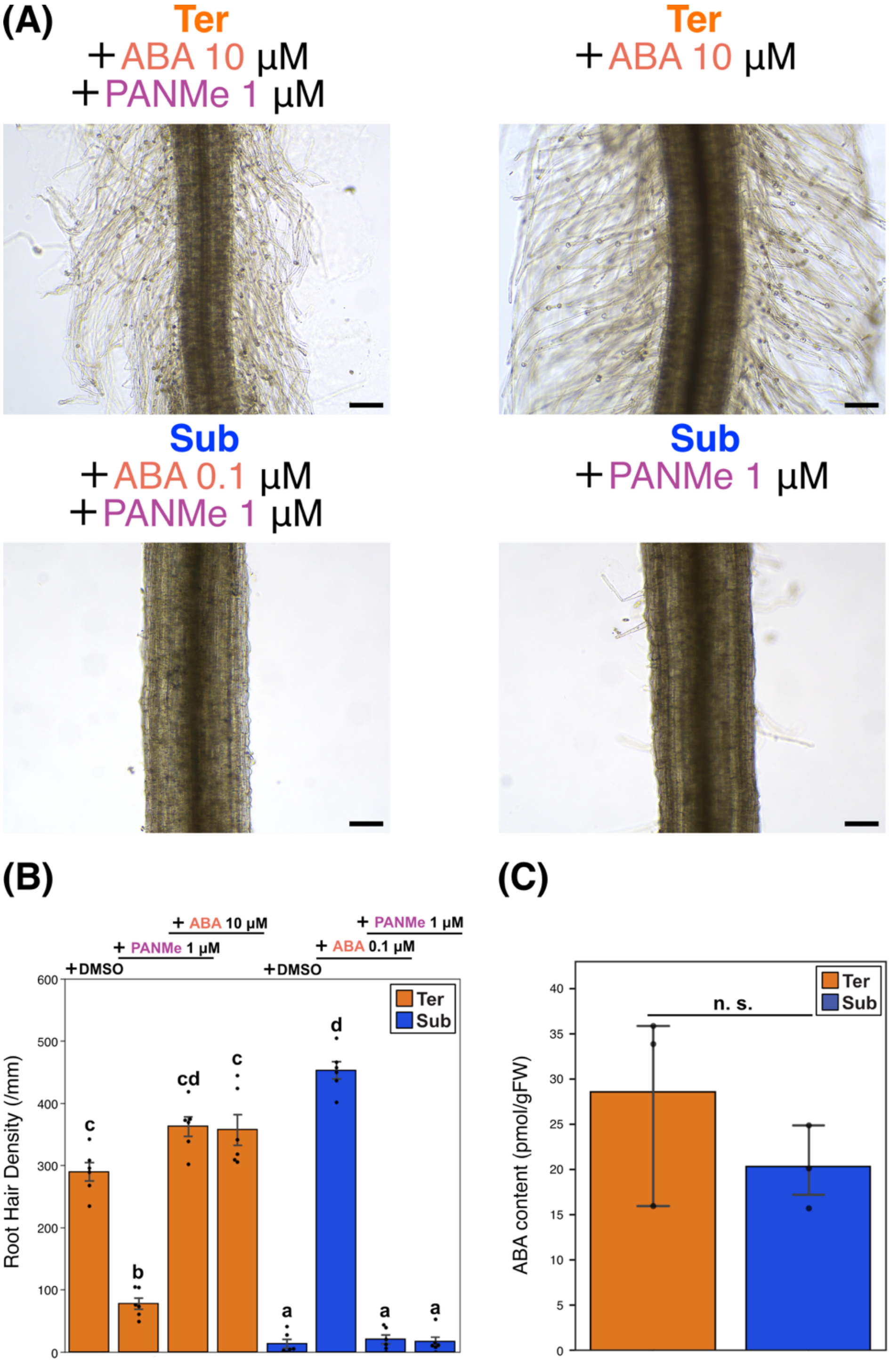


**Supplementary Figure 5.** Supplementary data of the effect of ABA on root hair development. **(A)** Effects of ABA and PANMe treatments, including single and combined applications, on root hair development. Scale bar: 100 µm. **(B)** Root hair density under the indicated treatments (n = 6). Error bars indicate the standard error of the mean (SEM). Different letters indicate statistically significant differences among treatment groups according to the Games–Howell test ($p < 0.05$). **(C)** ABA content under two conditions ($n = 3$). n.s.: not significant (Welch's t-test).

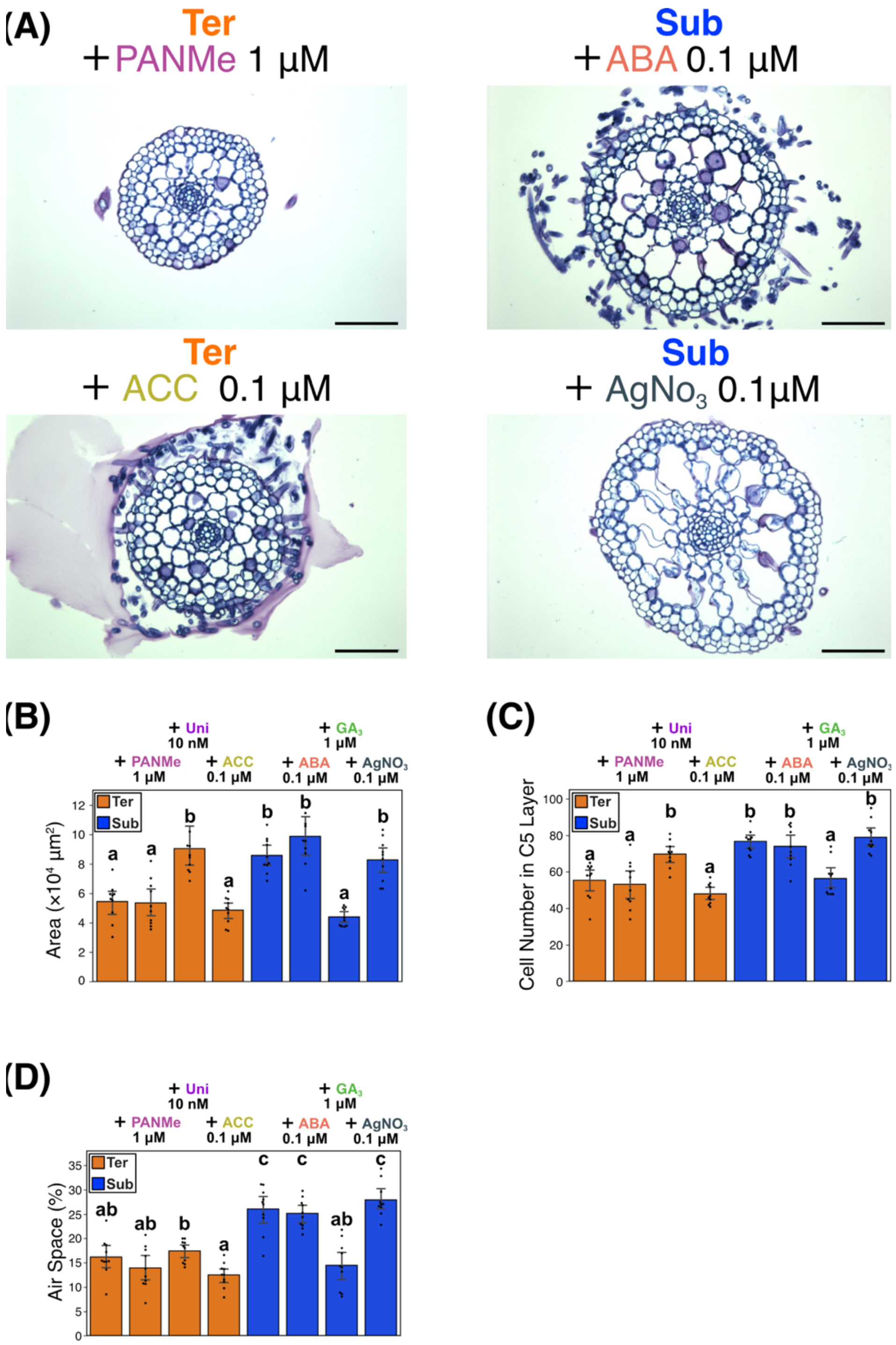


**Supplementary Figure 6.** Effect of other phytohormones on root anatomy. **(A)** Effects of ABA, ethylene, and their inhibitors on root anatomy. Scale bar: 100 µm. **(B–D)** Quantitative analyses of root anatomical traits. **(B)** Root cross-sectional area. **(C)** Epidermal cell number. **(D)** Proportion of air space. n = 6. Error bars indicate the standard error of the mean (SEM). Different letters indicate statistically significant differences among groups according to the Games–Howell test ($p < 0.05$).

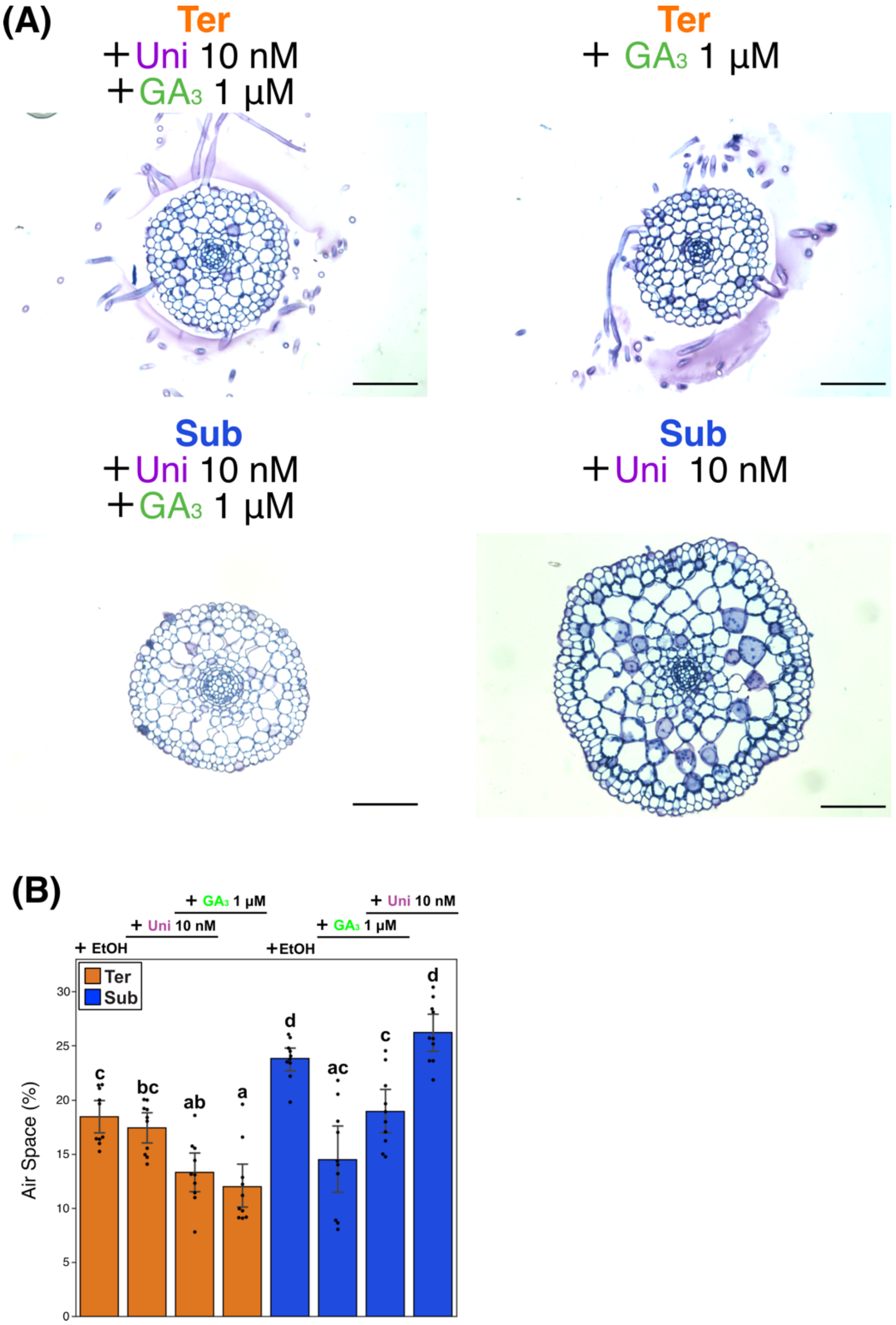


**Supplementary Figure 7.** Supplementary data of the effect of GA on root anatomy. **(A)** Effects of $GA_3$ and uniconazole P treatments, including single and combined applications, on root anatomy. Scale bar: 100 µm. **(B)** Proportion of air space under the treatments (n = 6). Error bars indicate the standard error of the mean (SEM). Different letters indicate statistically significant differences among treatment groups according to the Games–Howell test ($p < 0.05$).

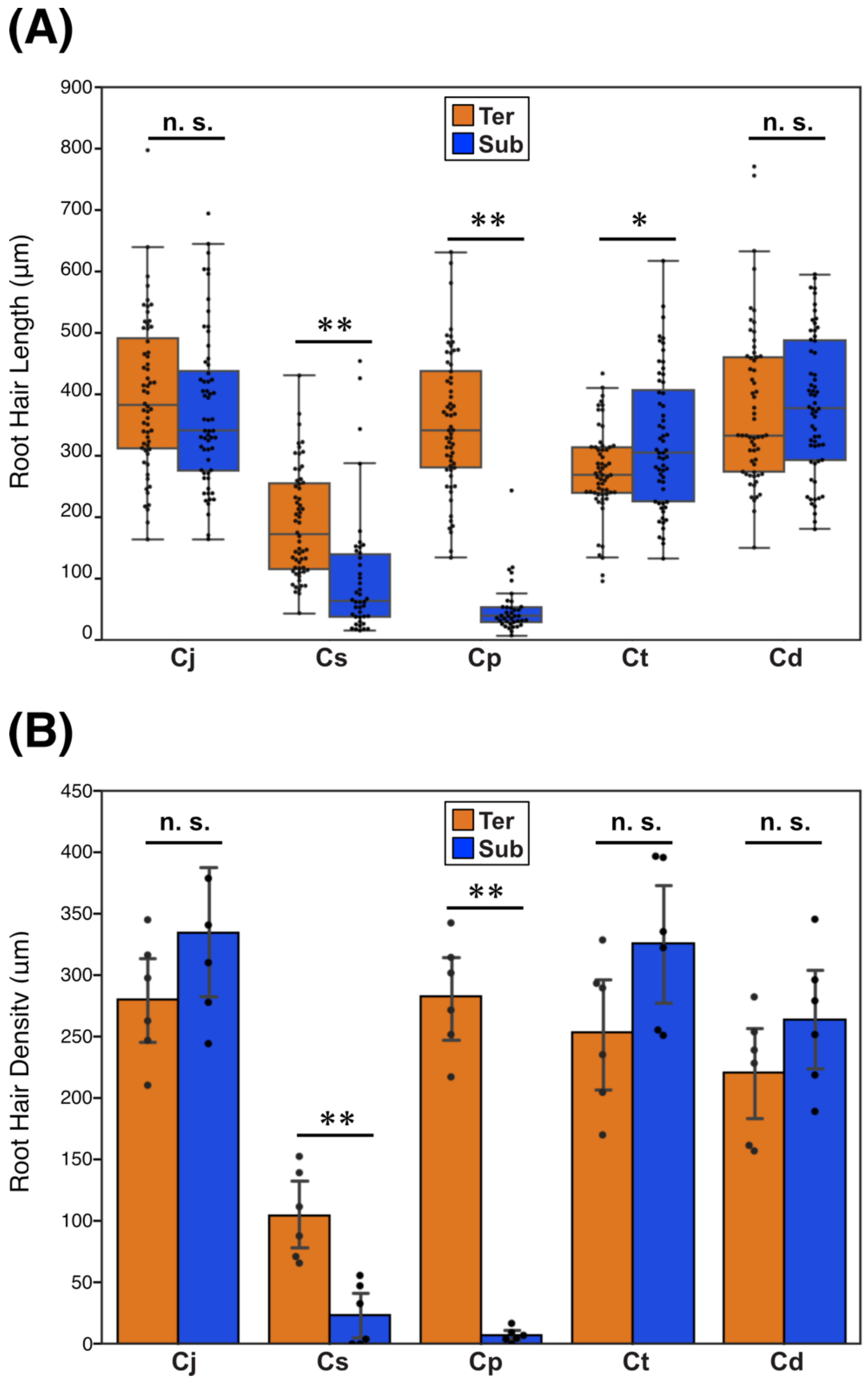


**Supplementary Figure 8.** Quantitative data of root hair traits of relative *Callitriche* species. **(A)** Difference of root hair length under two conditions. n = 38-60. **(B)** Difference of Root hair density under two conditions. n = 6. Error bars indicate the standard error of the mean (SEM). Asterisks indicate significant differences between conditions based on Welch's t-test followed by FDR (Benjamini–Hochberg) correction (n.s.: not statistically significant; $*$: $p < 0.05$; $**$: $p < 0.01$).

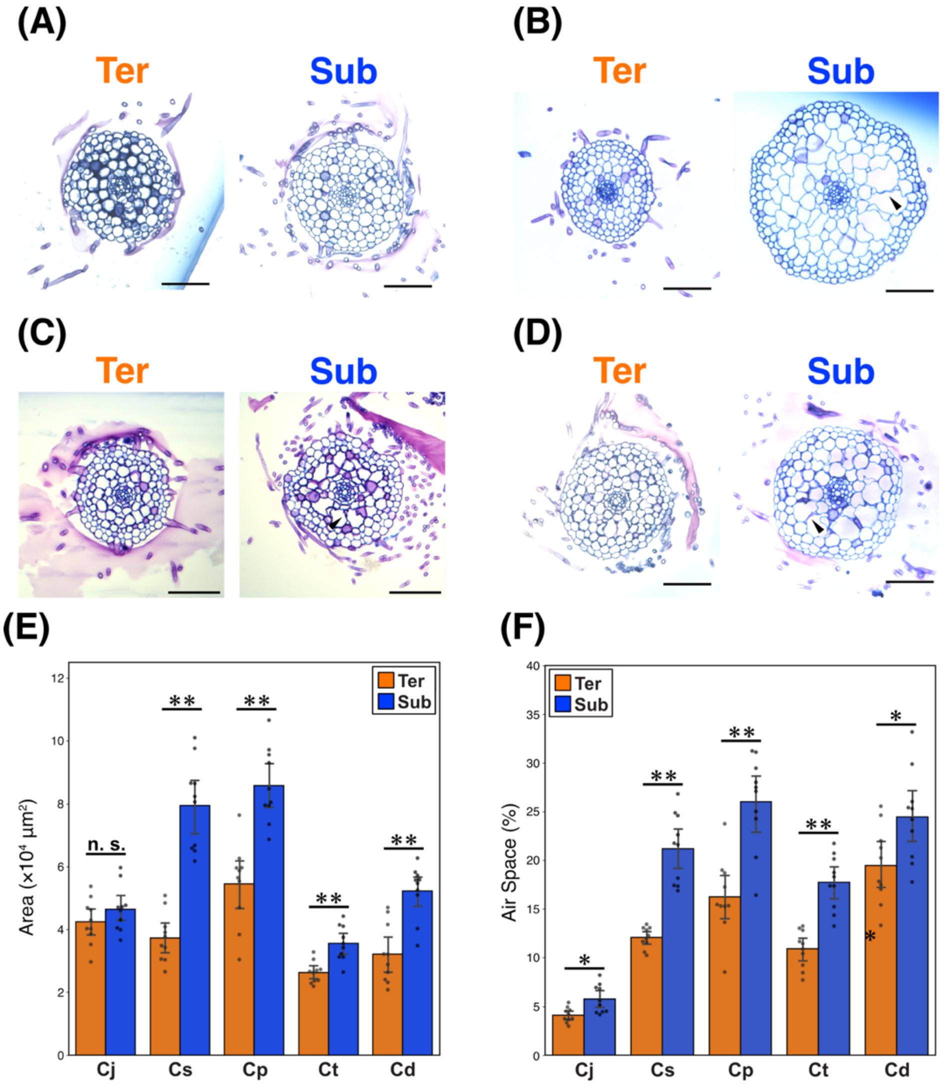


**Supplementary Figure 9.** Difference of the anatomy of relative *Callitriche* species. **(A-D)** Root cross-section of *Callitriche* species under two conditions. **(A)** *C. japonica*. **(B)** *C. stagnalis*. **(C)** *C. terrestris*. **(D)** *C. deflexa*. Arrowheads indicates dead cortical cells. Scale bar: 100 μm. **(E)** Difference of root cross-sectional area under two conditions. **(F)** Difference of the proportion of air space under two conditions. n = 10. Error bars indicate the standard error of the mean (SEM). Asterisks indicate significant differences between conditions based on Welch's t-test followed by FDR (Benjamini–Hochberg) correction (n.s.: not statistically significant; ∗: $p < 0.05$; ∗∗: $p < 0.01$).

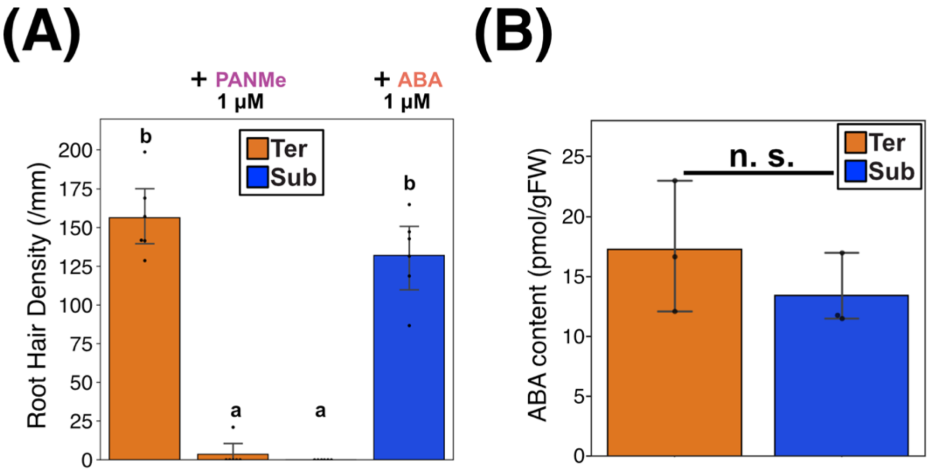


**Supplementary Figure 10.** Quantitative data of root hair density and endogenous ABA content in *L. arcuata*. **(A)** Root hair density under the indicated treatments (n = 6). Error bars indicate the standard error of the mean (SEM). Different letters indicate statistically significant differences among treatment groups according to the Games–Howell test (p < 0.05). **(B)** ABA content under two conditions (n = 3). n.s.: not significant (Welch's t-test).

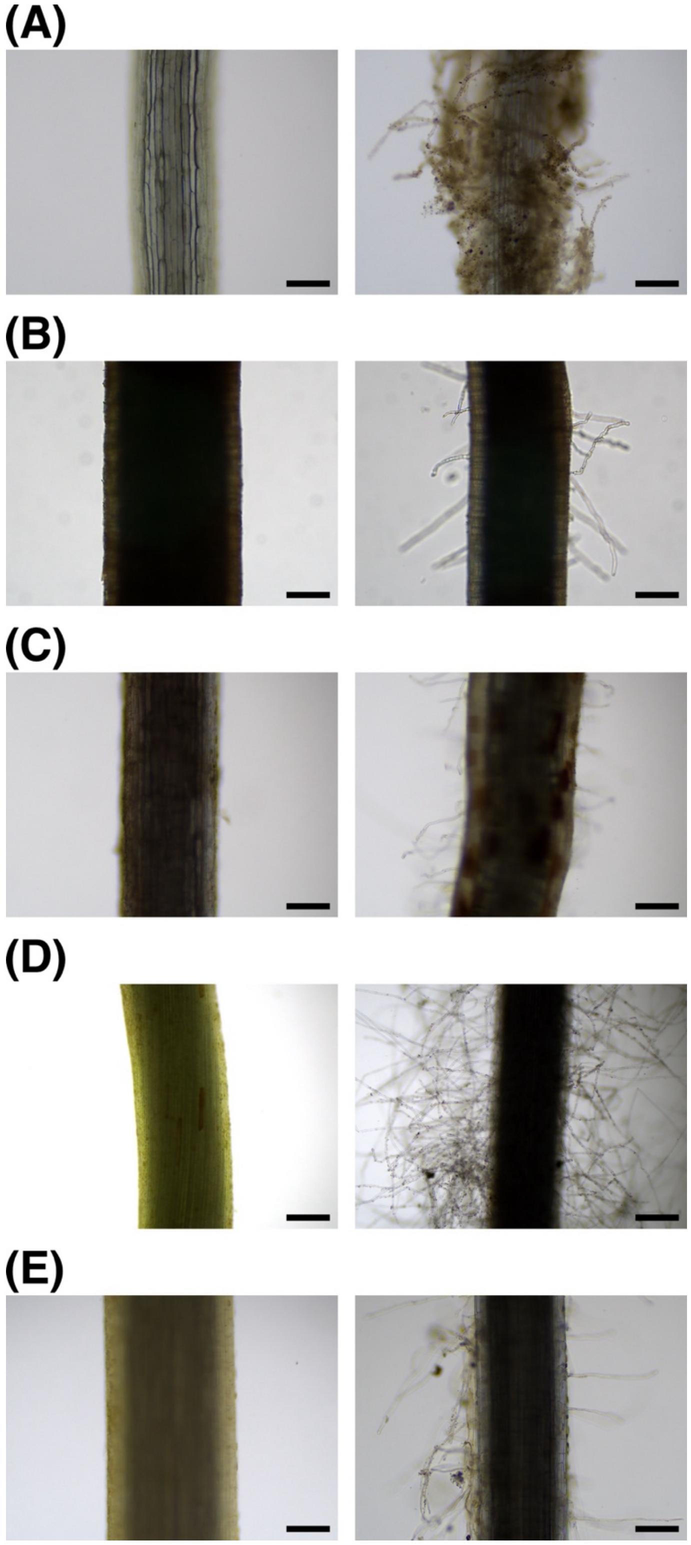


**Supplementary Figure 11.** Suppressed root hair development in water and its induction in soil in submerged aquatic plants. Representative images of roots from submerged aquatic plants. For each species, the same roots are shown in water (upper part of the root, left panels) and after entering the soil (lower part of the root, right panels). **(A)** *Callitriche palustris*. **(B)** *Ludwigia arcuata*. **(C)** *Rotala hippuris*. **(D)** *Egeria densa*. **(E)** *Potamogeton malaianus*. Scale bars: 200 µm.

**Supplementary Table 1. Sterile growth conditions for each species used in this study.**

| Species | Salt Mixture | Sucrose | Gellan Gum | pH | Source of materials |
|---|---|---|---|---|---|
| *Callitriche palustris* | ½ × MS | 2.0% (w/v) | 0.3% (w/v) | 5.8 | Collected in wild (Nagano, Japan) |
| *Callitriche deflexa* | ½ × MS | 2.0% (w/v) | 0.3% (w/v) | 5.8 | Collected in wild (Hyogo, Japan) |
| *Callitriche terrestris* | ½ × MS | 0.5% (w/v) | 0.3% (w/v) | 5.8 | Collected in wild (Nagano, Japan) |
| *Callitriche japonica* | 1 × B5 | 0.5% (w/v) | 0.3% (w/v) | 5.8 | Collected in wild (Ibaraki, Japan) |
| *Callitriche stagnalis* | ½ × B5 | 2.0% (w/v) | 0.3% (w/v) | 5.8 | Collected in wild (Tokyo and Kanagawa, Japan) |
| *Ludwigia arcuata* | ½ × B5 | 1.0% (w/v) | 0.3% (w/v) | 5.8 | Commercial aquarium supplier |

**Supplementary table 2. Information of root traits on non-*Callitriche* aquatic plants examined in this study.**

| Species name | Order | Class | Characteristics | Source of information | Source of material |
|---|---|---|---|---|---|
| *Potamogeton malaianus* | Alismatales | Potamogetonaceae | Few root hairs | Supplementary Fig. 10 | Collected in wild (Shiga, Japan) |
| *Egeria densa* | Alismatales | Hydrocharitaceae | Few root hairs | Supplementary Fig. 10 | Commercial aquarium supplier |
| *Elatine triandra* | Malpighiales | Elatinaceae | Cartwheel-like aerenchyma | Fig. 8 | Collected in wild (Kanagawa, Japan) |
| *Rorippa aquatica* | Brassicales | Brassicaceae | Cartwheel-like aerenchyma | Fig. 8 | Provided by Prof. Seisuke Kimura |
| *Ludwigia arcuata* | Myrtales | Onagraceae | Few root hairs, The plasticity of root hair development | Fig. 8 | Commercial aquarium supplier |
| *Rotala hippuris* | Myrtales | Lythraceae | Few root hairs | Supplementary Fig. 10 | Commercial aquarium supplier |
| *Myriophyllum spicatum* | Saxifragales | Haloragaceae | Few root hairs | Supplementary Fig. 10 | Collected in wild (Nagano, Japan) |
| *Hydrocotyle verticillata* var. *triradiata* | Apiales | Araliaceae | Cartwheel-like aerenchyma | Fig. 8 | Commercial aquarium supplier |

**Supplementary table 3. Information of aquatic plants examined in previous studies**.

| Species name | Order | Class | Characteristics | Source publications |
|---|---|---|---|---|
| *Eleocharis pusilla* | Poales | Cyperaceae | Absent or sparse root hairs | Crayton & Bagyaraj, 1984 |
| *Pontederia (Eichhornia) crassipes* | Commelinales | Pontederiaceae | No root hairs | Yin et al. 2025 |
| *Calla palustris* | Alismatales | Araceae | No root hairs | Shannon, 1953 |
| *Elodea canadensis* | Alismatales | Hydrocharitaceae | No root hairs in water | Cormack, 1937 |
| *Lemna minor* | Alismatales | Araceae | No root hairs | Shannon, 1953 |
| *Lemna trisulca* | Alismatales | Araceae | No root hairs | Shannon, 1953 |
| *Lemna yungenensis* | Alismatales | Araceae | Cartwheel-like aerenchyma | Ware et al. 2023 |
| *Orontium aquaticum* | Alismatales | Araceae | No root hairs | Shannon, 1953 |
| *Pistia stratiotes* | Alismatales | Araceae | No root hairs | Shannon, 1953 |
| *Spirodela intermedia* | Alismatales | Araceae | Cartwheel-like aerenchyma | Ware et al. 2023 |
| *Spirodela polyrhiza* | Alismatales | Araceae | No root hairs, Cartwheel-like aerenchyma | Shannon, 1953, Ware et al. 2023 |
| *Ranunculus rivularis* | Ranunculales | Ranunculaceae | Absent or sparse root hairs | Crayton & Bagyaraj, 1984 |
| *Ranunculus limosella* | Ranunculales | Ranunculaceae | Absent or sparse root hairs | Crayton & Bagyaraj, 1984 |
| *Elatine gratioloide* | Malpighiales | Elatinaceae | Absent or sparse root hairs | Crayton & Bagyaraj, 1984 |
| *Cardamine amara* | Brassicales | Brassicaceae | Cartwheel-like aerenchyma | Kudoh et al. 2025 |
| *Trapa natans* | Myrtales | Lythraceae | No root hairs | Shannon, 1953 |

| | | | | |
|---|---|---|---|---|
| *Myriophyllum triphyllum* | Saxifragales | Haloragaceae | Absent or sparse root hairs | Crayton & Bagyaraj, 1984 |
| *Myriophyllum propinquum* | Saxifragales | Haloragaceae | Absent or sparse root hairs | Crayton & Bagyaraj, 1984 |
| *Myriophyllum pedunculatum* | Saxifragales | Haloragaceae | Absent or sparse root hairs | Crayton & Bagyaraj, 1984 |
| *Crassula (Tillaea) sinclairii* | Saxifragales | Crassulaceae | Absent or sparse root hairs | Crayton & Bagyaraj, 1984 |
| *Lobelia (Pratia) perpusilla* | Asterales | Campanulaceae | Absent or sparse root hairs | Crayton & Bagyaraj, 1984 |
| *Lilaeopsis lacustris* | Apiales | Apiaceae | Absent or sparse root hairs | Crayton & Bagyaraj, 1984 |
| *Glossostigma elatinoides* | Lamiales | Phrymaceae | Absent or sparse root hairs | Crayton & Bagyaraj, 1984 |
| *Glossostigma submersum* | Lamiales | Phrymaceae | Absent or sparse root hairs | Crayton & Bagyaraj, 1984 |
| *Limosella lineata* | Lamiales | Scrophulariaceae | Absent or sparse root hairs | Crayton & Bagyaraj, 1984 |
